# Optimizing Lipidomics Analysis Workflows for Biological Fluids and Extracellular Vesicles with Integrated Liquid Chromatography Tandem Mass Spectrometry Approaches

**DOI:** 10.1101/2024.11.08.622653

**Authors:** Adriana F. L. Vilela, Miguel R. Patrício, Thiago V. Defelippo-Felippe, Viviani Nardini, Nathan N. H. Pontes, Jonatan C. Carvalho, Pedro Nobre-Azevedo, Daniel L. Rodrigues, Bianca T. M. Oliveira, Pedro V. da Silva-Neto, Ana P. M. Fernandes, Fausto Almeida, Lucia H. Faccioli, Carlos A. Sorgi

**Affiliations:** Departamento de Química, Faculdade de Filosofia Ciências e Letras de Ribeirão Preto – FFCLRP, Universidade de São Paulo – USP, Ribeirão Preto, 14040-901, SP, Brazil; Departamento de Bioquímica e Imunologia, Faculdade de Medicina de Ribeirão Preto – FMRP, Universidade de São Paulo – USP, Ribeirão Preto, SP, 14049-900, Brazil; Departamento de Análises Clínicas, Toxicológicas e Bromatológicas, Faculdade de Ciências Farmacêuticas de Ribeirão Preto FCFRP, Universidade de São Paulo – USP, Ribeirão Preto, 14040-903, SP, Brazil; Centro de Excelência em Quantificação e Identificação de Lipídios – CEQIL, Faculdade de Ciências Farmacêuticas de Ribeirão Preto – FCFRP, Universidade de São Paulo – USP, Ribeirão Preto, 14040-903, SP, Brazil; Departamento de Enfermagem Geral e Especializada, Escola de Enfermagem de Ribeirão Preto-EERP, Universidade de São Paulo-USP, Ribeirão Preto 14040-902, SP, Brazil

**Author notes:** Corresponding author: Carlos A. Sorgi, email adress, telefone number: /8871, affiliation address: Departamento de Química, Faculdade de Filosofia Ciências e Letras de Ribeirão Preto, Av. Bandeirantes, 3900, 14040-901, Ribeirão Preto, SP, Brazil. These authors contributed equally to this work and share first authorship (Adriana F. L. Vilela, Miguel R. Patrício).

**Keywords:** Lipidomics, extracellular vesicles, lipid-extraction, LC-MS/MS, Bioinformatics

## Abstract

Lipidomics, a subfield of metabolomics, involves the comprehensive analysis of lipids within biological systems and has become a cornerstone of biomedical research, driven by recent technological advancements. Lipids are crucial biomolecules in cellular functions and have been increasingly recognized for their roles in physiological and pathological processes. This study focuses on innovative strategies for developing, validating, and applying comprehensive analytical methods for untargeted lipidomics using liquid chromatography-tandem mass spectrometry (LC-MS/MS) in human plasma and extracellular vesicles (EVs). We describe improvements based on analytical validation parameters, including inter-day repeatability, limit of quantification, precision, accuracy, recovery, and matrix effects. Plasma samples were used as a proof-of-concept study, and the method was ultimately applied to human macrophage-derived EVs. Samples preparations were achieved through four liquid-liquid extraction methods for lipids in order to achieve a broad coverage of lipid classes as well as high recovery and repeatability. Additionally, we demonstrated that a sonication-assisted homogenization step effectively facilitates lipid extraction from EVs. Through untargeted lipidomics, our study identifies and quantifies a diverse range of lipid species in human plasma (225 molecular lipids) and macrophage-derived EVs (124 molecular lipids) within different classes. Overall, we present an innovative methodology that combines pre-analytical lipid extraction techniques with high-resolution LC-MS/MS to enhance lipidomics research. This approach holds promise for personalized medicine and the discovery of novel lipid cargo associated with the various biological pathways involved with EVs biogenesis.

## 1. Introduction

Lipidomics, the intricate analysis of lipids within biological systems, has recently seen a wave of innovation, following new possibilities in biomedical research. The lipids are a varied class of biomolecules, crucial in cellular functions, acting as structural parts of cell membranes, energy stores, and signaling molecules [1,2]. Beyond these basic roles, lipids are increasingly recognized for their involvement in many physiological and pathological processes, including cardiovascular disease, cancer, and neurodegenerative disorders [3,4]. Advances in liquid chromatography-mass spectrometry (LC-MS) have been essential to transforming lipidomics, enabling the identification and quantification of an ever-growing range of bioactive lipids [1]. Specifically, ambient ionization techniques have enhanced mass spectrometry’s capabilities, allowing for rapid, real-time, *in situ*, and *in vivo* analysis of lipid profiles in complex biological samples [5]. Today, lipidomics is routinely used in discovering biomarkers, detecting early diseases, and understanding lipid-mediated signaling pathways, aiding the creation of new therapeutic strategies and personalized medical approaches.

Cellular and tissue lipids encompass a vast array of unique molecular structures, comprising hundreds of thousands of distinct lipid species. An actual comprehensive classification system grouped individual lipid molecules into eight main categories: fatty acids, glycerolipids, glycerophospholipids, sphingolipids, sterol lipids, prenol lipids, and saccharolipids [6]. Within each category, lipid species are further subdivided into distinct lipid classes based on their polar head group structures. For instance, glycerophospholipids derive from phosphatidic acids (PA) are categorized into choline (PC), ethanolamine (PE), inositol (PI), serine (PS) and glycerol (PG) glycerophospholipid, depending on the phospho-head groups, respectively, attached to a glycerol backbone with fatty acyl chains [7]. This structural diversity of lipids has important implications for their biological functions. Non-polar lipids, like triglycerides (TG), serve as energy storage; while polar lipids, like phospholipids, are the major structural components of cell membranes, contributing to their physical properties and providing scaffolding for membrane proteins [8]. Moreover, numerous lipid subclasses and individual molecular species function as signaling molecules, regulating critical cellular processes such as cell proliferation, inflammation, and apoptosis [9–11]. The comprehensive and quantitative analysis of these lipid structures is challenging.

Extracellular vesicles (EVs) are released by nearly all cell types and are found in various biological fluids, making them a key element of cellular signaling. Several studies have highlighted the involvement of EVs, which carry diverse types of biomolecules cargo, including RNA, lipids, and proteins, in both physiological functions and pathological states such as inflammation and cancer [12]. Although most works have extensively analyzed the protein and microRNA content of EVs, our knowledge about the lipid components remains limited, despite their crucial roles in EV formation, release, targeting, and cellular uptake. EV membranes often show an accumulation of cholesterol, sphingolipids, and saturated phospholipids[13] . Since EVs from different biological fluids reflect the molecular profiles of their originating cells, they hold potential as non-invasive biomarkers for diagnosing various diseases.

In this way, exploring new techniques and strategies to enhance the accuracy and reproducibility of lipid analysis is mandatory [1,14]. One critical aspect of lipidomics research is the pre-analytical phase, which encompasses the sample preparation steps prior to mass spectrometric analysis. Advances in lipid extraction methods have played a pivotal role in addressing the unique challenges associated with the analysis of complex biological samples [15]. Traditional lipid extraction techniques, such as reported by Bligh & Dyer (1959), have been widely used, but they often suffer from incomplete extraction, poor reproducibility and the potential loss of certain lipid species [16]. To overcome these limitations, alternative pre-analytical strategies that aim to improve the recovery, selectivity, and quantitative accuracy of lipid extraction were introduced and explored herein [15]. In parallel, the development of targeted and non-targeted lipidomics by high-resolution LC-MS/MS approaches has further expanded the capabilities of lipid analysis [1]. Targeted lipidomics, which focuses on the quantification of specific lipid species, has demonstrated improved accuracy and precision; while non-targeted lipidomics has enabled the discovery of novel lipid biomarkers and the investigation of uncovered lipid metabolic pathways [1].

The combination of the advancements in ionization techniques and high-resolution LC-MS/MS lipidomics strategies has significantly contributed to the improved accuracy and reproducibility of lipidomics research, particularly in the analysis of complex biological matrices [1,5,15]. In the present study, we performed sample preparation employing the most widely used liquid-liquid extractions methods in a human plasma and EVs matrixes. Furthermore, we established best analytical practices to validate our LC-MS/MS methodology for this lipidomics approach.

## 2. Experimental

### 2.1 Material and reagents

Methanol, acetonitrile, isopropanol, chloroform, hexane (all HPLC-grade), methyl tert-butyl ether (MTBE, 99.8%), formic acid (98%), acetic acid (99%), ammonium formate and ammonium acetate were purchased from Sigma-Aldrich (St. Louis, MO, USA). Ultrapure water used in all the preparations was obtained from a MILLI-Q® system (Millipore, USA). Synthetic lipid standards were purchased from Avanti Polar Lipids, Inc. (Alabaster, AL, USA), and are listed in Table S-1. Dulbecco’s Modification of Eagle’s Medium (DMEM) was acquired by Corning® (Arizona, USA); phorbol 12-myristate 13-acetate (PMA) and Lipopolysaccharide (LPS) by Sigma-Aldrich (St. Louis, MO, USA); fetal bovine serum (FBS) and Antibiotic Antimycotic Solution (100×) were acquired by Thermo Fisher Scientific Inc. (Waltham, MA, USA). The cytokines, such as IFN-γ, IL-4 and IL-13 were obtained by R&D Systems (Minneapolis, MI, USA).

### 2.2. Biological Samples

#### 2.2.1 Human plasma samples

Human plasma samples from healthy individuals were obtained from “PLURAL clinical study” of the Universidade de São Paulo (USP). This study strictly adhered to ethical principles and regulatory norms, as outlined in the Certificate of Presentation for Ethical Consideration (CAAE) - Process CAAE# 30525920.7.0000.5403. All subjects provided informed consent. Blood samples were collected in EDTA tubes and immediately put on ice for 30 min, then samples were centrifuged for 5 min, 5,000 g at 4°C. Plasma was collected, divided in 250 μL aliquots, added one volume of MeOH and storage at −80 °C until further analysis.

#### 2.2.2 Extracellular Vesicles (EVs)

##### 2.2.2.1 Macrophage Culture

The human acute monocytic leukemia (THP-1) cells were cultured at 37°C in a humidified atmosphere of 5% CO_2_ in DMEM medium containing 4.5 g/L glucose, L-glutamine and sodium pyruvate, supplemented with 500 UI penicillin, 500 µg streptomycin, 1.25 µg amphotericin and 10% FBS heat-inactivated and depleted of EVs (DMEM-c) [1]. Additionally, THP-1 cells (2.0 x 10^5^/mL) were cultured in DMEM-c with 25 nM Forbol 12-miristato-13-acetato (PMA) for 24 h at 37°C, for adherence and to obtain phenotype naïve macrophages. Macrophages were then washed once with DMEM to remove non-adherent cells and the remaining cells were used for the polarization protocol *in vitro* with the presence of IFN-γ (20 ng/mL) + LPS (1 µM) or IL-4 and IL-13 (1 µM) diluted in DMEM-c for 24 h at 37°C.

##### 2.2.2.2 EVs Isolation and purification

The required volume (40 mL) of macrophage culture supernatants were sequential centrifuged at room temperature by 5,000 g, 15 min and 15,000 g, 15 min. The supernatants obtained were concentrated in an Amicon system (Millipore, MA, USA) with molecular exclusion of 100 kDa. The concentrated supernatant was filtered at 0.45 µm and ultracentrifuged at 100,000 g, 1 h at 4°C. Then, the pellet obtained was re-suspended in 1 mL of water for subsequent analysis and/or stored at -80 °C. The purity, concentration, and particle size of those EVs samples were carried out using the Nanosight NS300 nanoparticle tracking analysis (NTA) function (Malvern Instruments, Malvern, UK). EVs pooled sample was prepared by mixing EV-derived from naive and polarized macrophages (equal proportions) at final concentration of 1 x 10^10^ particles/mL. Freshly isolated EVs were subjected, or not, to low-power sonication-treatments previously to the extraction assay. The ultrasonication bath operating parameters include bath temperature (25 °C), time cycle (0, 5, 10, 25 or 60s), and emitted ultrasound properties by 100 W with a frequency of 40 kHz (SolidSteel - SSBu, SP, Brazil).

### 2.3. Untargeted lipidomics workflow

#### 2.3.1 Standard solutions

Synthetic lipid standards (Table S1): monoacylglycerosphosphocholine (LPC) 17:0, monoacylglycerosphosphate (LPA) 17:0, diacylglycerophosphocholine (PC) 17:0/17:0, diacylglycerophosphatidylserine (PS) 17:0/17:0, diacylglycerophosphoethanolamine (PE) 17:0/17:0, diacylglycerophosphoglycerol (PG) 17:0/17:0), sphingomyelin (SM) 12:0 and ceramide (Cer) 17:0, were used as internal lipids standard (IS) for this work, representing the principal classes of lipids in cell membrane. Stock solutions of each synthetic IS were prepared by dissolving in chloroform/MEOH (9:1 or 5:1, *v/v*) at concentrations ranging from 25 mM to 5 mM and stored at -80 °C. Working solutions were prepared by successive dilution of stock solutions. A serial dilution was performed in the following concentrations: 20.0, 15.0, 12.5, 10.0, 7.5, 5.0, 2.5, 1.0, 0.5, 0.25 µM for the calibration curve. To optimize the compound parameters (collision energy [CE] and declustering potential [DP]), a standard solution of each IS was prepared at 10 µM in methanol acidified with formic acid (0.1%).

#### 2.3.2. Biological Samples extractions

Extraction of lipids from plasma or EVs samples was carried out based on the physico-chemical properties of the classical Matyash [17] and Bligh & Dyer [18] protocols, with some optimization modifications. Briefly, four liquid-liquid extractions methods were evaluated in our pre-analytical approach, such as MTBE-extraction protocol (MT), MTBE-extraction protocol modified (MM), Chloroform-extraction protocol (BD) and a hybrid MTBE and Chloroform sequential-extraction protocol (HD). Extractions were conducted with all the solvents on ice and the samples were prepared in quintuplicate. All the samples were spiked with 40 µL mixing of IS. The samples were mixed, incubated and centrifuged as described individually below. At the end of extractions, all collected samples were dry in vacuum-systems (SpeedVac SPD 1030, Thermo Fisher Scientific, CA, USA) pressure of 10.0 Torr at 45°C. The dried samples were re-suspended in 40 µL isopropanol/ acetonitrile/ H_2_O (2:1:1 *v/v/v*) and analyzed by LC-MS/MS system.

##### 2.3.2.1. MTBE-extraction protocol (MT)

Plasma (200 μL) or EVs (100 μL - 1 x 10^9^ particles) samples in glass tubes were adjusted volume with deionized ultrapure-water, 50 μL (for plasma) and 150 μL (for EVs), mixed with 260 μL MeOH. Followed by addition of MTBE (1 mL) and vortexed for 10s. Then, the tubes were put on a shaker for 1 hour, under gentle stirring at room temperature. Afterwards, the extraction solution was centrifuged at 10,000 g for 10 min followed by the collection of the upper organic-phase.

##### 2.3.2.2. MTBE-extraction modified protocol (MM)

Plasma (200 μL) or EVs (100 μL - 1 x 10^9^ particles) samples in glass tubes were adjusted volume with deionized ultrapure-water, 160 μL (for plasma) and 260 μL (for EVs), and mixed with 160 μL methanol. Followed by the addition of 540 μL MTBE and vortexed for 10s. The tubes were put on a shaker for 25 min, under gentle stirring at room temperature. Afterwards, the extraction solution was centrifuged at 10,000 g for 10 min followed by the collection of the upper organic-phase.

##### 2.3.2.3. Chloroform-extraction protocol (BD)

Plasma (200 μL) or EVs (100 μL - 1 x 10^9^ particles) samples were mixed with 710 μL of chloroform/MeOH (1:2 *v/v*) and 40 μL IS in glass tubes. Deionized ultrapure-water (100 μL) was added for adjusted volume in EVs sample tube. The tubes were vortexed and centrifuged at 10,000 g for 5 min, 4°C. The samples were followed by the addition of 250 μL chloroform and 250 μL deionized water and well vortexed, centrifuged at 10,000 g for 10 min to induce phase separation. Then, the lower organic phase (chloroform) was collected and stored. Afterwards, the re-extraction was carried out by addition of 500 μL chloroform to the aqueous phase, the mixture obtained was again well vortexed and centrifuged at 10,000 g for 10 min. The lower phase was collected and stored in the same tube as the previous extraction.

##### 2.3.2.4. Hybrid-extraction protocol (HD)

Plasma (200 μL) or EVs (100 μL - 1 x 10^9^ particles) samples were mixed with 710 μL of chloroform/MeOH (1:2 *v/v*) and 40 μL IS in glass tubes. Deionized ultrapure-water (100 μL) was added for adjusted volume in EVs sample tube. The tubes were vortexed and centrifuged at 10,000 g for 5 min, 4°C. The samples were followed by the addition of 250 μL chloroform and 250 μL deionized water and well vortexed, centrifuged at 10,000 g for 10 min to induce phase separation. Then, the lower organic phase (chloroform) was collected and stored. The sequential step was the re-extraction by adding 1 mL of MTBE to the aqueous phase of the remaining tube, put on a shaker for 10 min, under gentle stirring at room temperature. Then, centrifuged at 10,000 g for 10 min, 4°C. The organic phase was collected and added to the same tube of the previously chloroform extraction.

#### 2.3.3. Tandem mass spectrometry analysis

LC-MS/MS analyses were performed using ultra-high-performance liquid chromatographic (UHPLC - Nexera X2; Shimadzu, Kyoto, Honsh, Japa) coupled to a triple quadrupole time-of-flight (TripleTOF^®^ 5600^+-^Sciex, Foster, CA, USA) mass spectrometer. The UHPLC system consists of two LC 30AD pumps, an autosampler (SIL-30AC), a CTO-30A oven, a CBM-20A controller and DGU-20A degassing. The TripleTOF^®^ 5600^+^ mass spectrometer was equipped with a turbo-V IonSpray and calibrant delivery system (CDS). Data acquisition was accomplished on a Shimadzu CBM-20A system interfaced with a computer, using the Analyst^®^ TF software version 1.7.1 (SCIEX, CA, USA). The chromatographic separation was carried out using an Acquity UPLC® CSH™ C18 column (100 × 2.1 mm; 1.7 μm) from Waters (Milford, USA). The mobile phases used were: phase A (0.1% formic acid and 10 mM of ammonium formate in water with acetonitrile - 39.9:60 *v/v*) and phase B (0.1% formic acid and 10 mM of ammonium formate in isopropanol with acetonitrile - 89.9:10 *v/v*). Column oven was set at 45°C. Separations were made with a gradient as follows: 0.01-3 min, 30% B; 3.1-5.1 min, 43% B; 5.21-15.21 min, 65% B; 15.22-21.22 min, 85% B; 21.23-23.5 min, 100% B. The injection volume was 5 μL and the flow rate was set at 0.4 mL.min^-1^. Mass spectra were acquired from mass to charge ratio (*m/z*) 50 to 1250. The electrospray ionization (ESI) source operated in positive and negative mode. MS conditions were as follow: nebulizer gas (GS1) at 50 psi, turbo gas (GS2) at 60 psi, curtain gas (CUR) at 25 psi, electrospray voltage (ISVF) at +4,500 V and −4,500V, and turbo ion spray source temperature at 500°C. The dwell time was set at 10 ms, and a mass resolution of 35,000 was achieved at *m/z* 400. External calibrations of the calibrated delivery system (CDS) were carried out using an atmospheric-pressure chemical ionization probe (APCI). Automatic mass calibration (<1 ppm) was performed after each of the five sample injections using APCI Positive and negative Calibration Solution (SCIEX, CA, USA) injected via direct infusion at a flow rate of 350 μL/min.

#### 2.3.4. Method validation

The analytical method was validated according to the Food and Drug Administration (FDA) guidelines [19]. The parameters, such as linearity, intra- and inter-day precision, accuracy and the lower-limits of quantification (LLOQ) were evaluated. Linearity and lower limit of quantification (LLOQ) were assessed using the external calibration curve of an IS lipid mixture at concentrations range of 0.25–25 µM. Calibration curve and quality control (QC) samples were prepared by serial dilution as described in section 2.3.1. The solutions were prepared in triplicate, mixed for 10 s, and 5 µL aliquots were injected into the LC-MS/MS system. Calibration curves were constructed by plotting the standard area against its nominal concentration and the linear equations were obtained from regression analysis by the least-squares method. Acceptance criteria were set to a deviation of ≤15% from the nominal value for concentrations. The coefficient of regression (r2) should always be > 0.97 for acceptance. LOQs were evaluated by considering a signal-to-noise (S/N) ratio ≥ 10 and RSD ≤ 20%. Intra- and inter-day precision and the method accuracy were evaluable by injections of three different concentrations of lipids mixture standards, named quality controls (QCs): 3µM for low-concentration (LQC), 10µM for medium-concentration (MQC), and 20µM for high-concentration (HQC). Five samples of each concentration were prepared and analyzed on the same day and three non-consecutive days. According to FDA guidelines, to be validated, the relative standard deviation (RSD%) must be less than 20 % and the accuracy and repeatability must be between 80 and 120 %. The method selectivity was evaluated using blank runs in which the target analytes are not present and measured under identical conditions.

##### 2.3.4.1 Recovery and matrix effects

The efficiency of extractions for the lipids analytes in the biological matrices was determined. For this, the recovery was assessed by using IS solutions. Plasma samples were spiked with IS before extraction, and the matrix effects were evaluated. Recovery was determined by comparing the analytical response of IS (at 25 µM) for the extracted samples to the IS quantification value directly prepared in methanol. The ratio between mean peak areas of IS spiked samples extraction and the standard samples represent the recoveries for each analyte. Recovery was expressed as the percentage of the expected value, as calculated by the average area ratio between the extraction samples and the standard samples. Matrix-dependent effect was established using the peak areas after extraction compared with the corresponding peak areas of the pure IS solutions. Samples for matrix effect and recovery studies were prepared in quintuplicate, and a CV of ≤20% was used as the accepted criteria for QC.

### 2.4. Bioinformatics and statistical analyses

All datasets acquired were analyzed using the softwares PeakView 2.1 (Sciex, CA, USA), MultiQuant^TM^ 3.0.2 (Sciex, CA, USA) and MS-Dial^TM^ 5.1.230429 (open-source software -RIKEN, Yokohama, Kanagawa, Japan). Targeted data, for method validation, was quantified by the Sciex softwares and the untargeted data were extracted and aligned using MS-Dial. Untargeted datasets raw files into .wiff format were extracted using Analyst^®^ TF (version 1.7.1.) software to initiate the process in MS Dial software. The setting parameters used are described as follows: i) raw measurements files were type SWATH acquisition; ii) the measurements parameters to MS/MS were centroid data for the negative and positive modes; iii) data collection was obtained as set; *m/z* tolerance to MS2 at 0.05 Da and mass range of 50-1250; iv) the parameters to peak detection included the following: 50% of amplitude, mass slice width of 0.1 Da, linear weighted moving average, smoothing level and minimum peak width of 5 and 3 scans, respectively; v) spectrum deconvolution parameters were sigma window value of 0.5 and MS/MS abundance cut off of 90 amplitude; vi) for the annotation lipids was used MS-DIAL -*TandemMAssSpectralAtlas-VS69* file as reference library (http://prime.psc.riken.jp/compms/msdial/main.html#MSP). The parameters included: accurate mass tolerance of 0.05 Da; dot product score, 600; weighted dot product score, 600; reverse dot product score, 800; matched spectrum percentage, 70%; minimum number of matched spectrum; vii) features coming from multiples adducts ([M+Na]^+^, [M+K]^+^, [M+NH_4_]^+^, [M+NaCOOH]^+^ in positive mode, [M+HCOOH]^−^ and [M+CH_3_COOH]^−^ in negative mode) were filtered by retaining in the final data set only the highest MS intensity signal from each cluster; viii) alignment parameters were used reference file as control without matrix, retention time tolerance set as 0.1min and MS2 tolerance at 0.05 min. Finally, all the structures of the annotated lipids were confirmed by matching structural fragments with MS/MS experimental fragmentation available in MS-Dial reference library. The resulting data set was exported in .svg or .xls format and is ready for further data analysis. Statistical analysis and graphics data was performed using GraphPad Prism software version 9.0 (Prism, LaJolla, CA), the group graphics data were expressed as the means ± s.e.m. and statistical significance was set at *p* < 0.05. Partial Least Squares Discriminant Analysis (PLS-DA) and heatmaps were determined using the MetaboAnalyst 6.0 online software (https://www.metaboanalyst.ca/).

## 3. Results and Discussion

### 3.1 Tandem mass spectrometry analysis optimization parameters

Lipids encompass a vast array of molecules with diverse physicochemical properties. For the broad identification of lipid species from biological matrices, several analytical steps must be carefully optimized, including extraction, chromatographic separation, and mass spectrometry analysis. The goal of untargeted, high-resolution mass spectrometry-based lipidomics is to identify as many individual molecular species as possible within the sample. In this context, we developed and validated a label-free untargeted lipidomics approach for application to human plasma and macrophage-derived EVs. The internal standards used for method validation included commonly employed lipid species in lipidomics, such as phospholipids (phosphatidylcholines, phosphatidylethanolamines, phosphatidylserines, phosphatidylglycerols), lysophospholipids and sphingolipids (sphingomyelin, and ceramides). LC-MS/MS method was developed on the basis of reported protocols in literature [20–22]. To enhance coverage of lipid classes, particularly nonpolar ones such as TG, we extended analysis times to 25 min. Liquid-chromatographic separation was based in reversed phase-LC (RP-LC) and the elution gradient of chromatographic conditions sought to identify multiple classes of polar and nonpolar lipids. All the lipids IS used were identified and quantified in negative and positive ionization, as shown in Table S2.

Extracted-ion chromatograms (XIC) and fragmentation profile of lipids analyzed simultaneously are demonstrated in Fig. 1 A-B. Established on the best ionization mode and peak intensity, positive ionization was used for the species LPC, that was eluted first, followed by SM, PS, PC; negative ionization was used for LPA, PG, Cer and PE, in order to the retention time in RP-LC analysis. The separation mechanism in RP-LC of lipids is based on lipophilicity, thus, lipid species containing longer acyl-chains are eluted from the LC column later than shorter chain lipids, and saturated acyl structures are eluted later than polyunsaturated analogs.

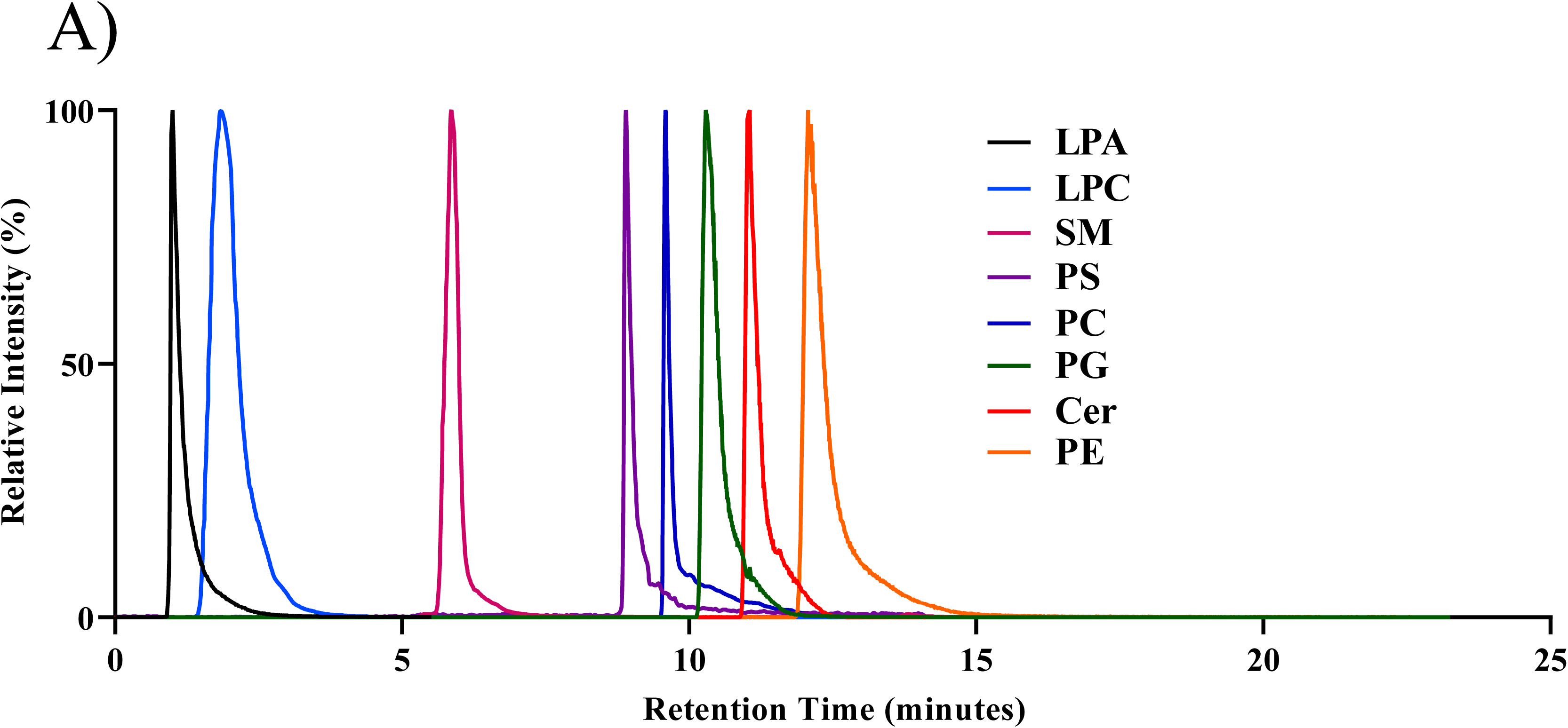
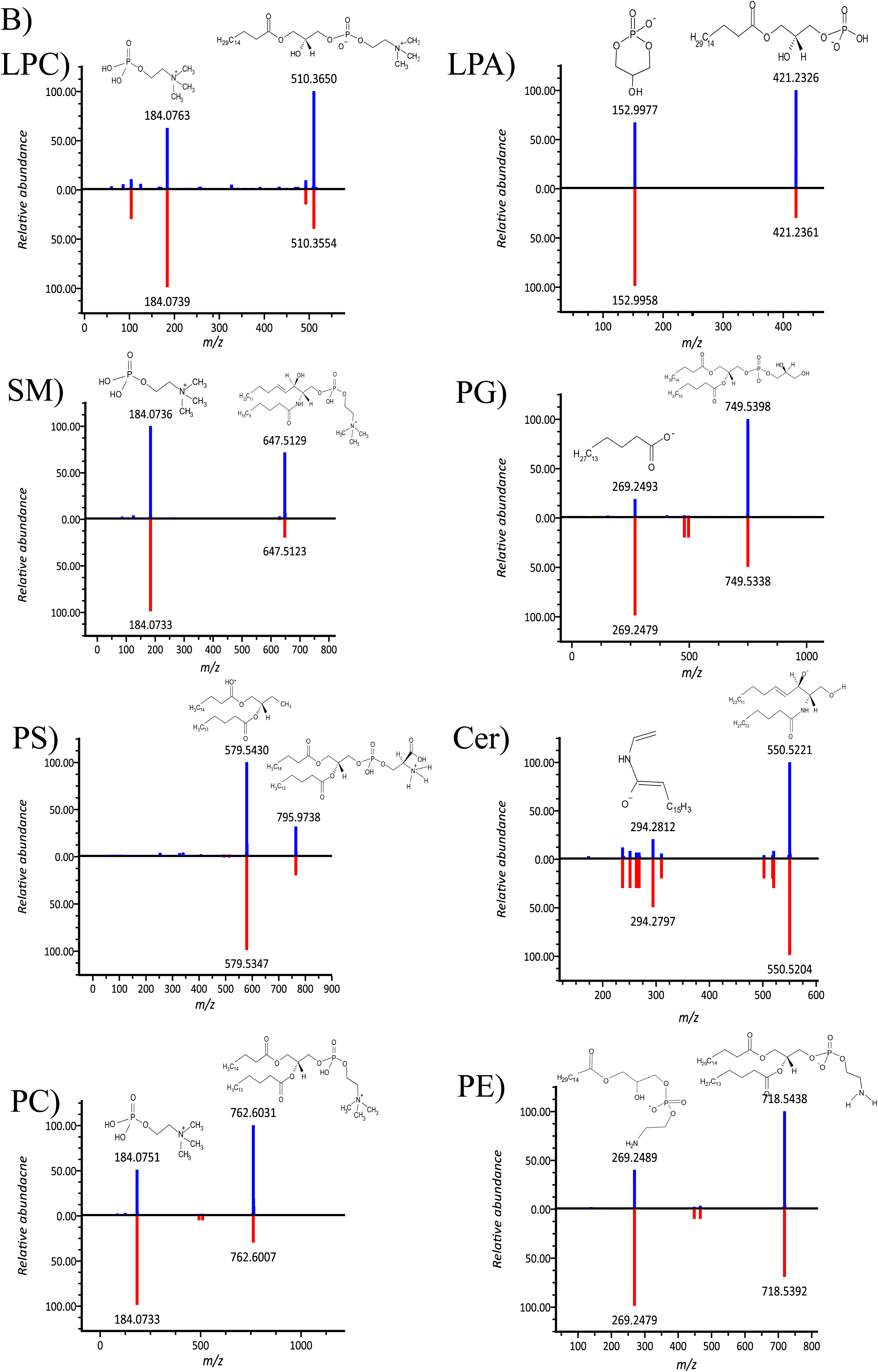

The ionization mode employed in LC-MS/MS analysis is a critical factor that significantly impacts the detected lipid profile [1,5]. Specifically, when utilizing electrospray ionization (ESI) as the ion source, certain lipid classes demonstrate preferential ionization in the positive mode, as showed for PC, LPC, and SM, while others, such as PI, PS, and phosphatidic acid (PA), are better ionized in the negative mode [1,23]. In this work, the optimal ionization mode for each lipid standard was determined based on the largest signal-to-noise ratio, with the exception of lysophosphatidic acid (LPA), which was analyzed in the negative mode, and lysophosphatidylcholine (LPC), which was analyzed in the positive mode (Table S2). The ability to selectively target lipids of interest through the appropriate choice of ionization mode is a crucial aspect of comprehensive lipidomics analysis, as it allows for the improved detection and quantification of a diverse range of lipid species within complex biological matrices [5,23]. Indeed, our results demonstrated that the LC-MS/MS method used suitably for the identification of different lipid classes and accurately quantified.

### 3.2 Untargeted lipidomics method validation

The validation of an analytical method is crucial for ensuring the reproducibility of results, particularly when dealing with sensitive and complex techniques such as LC-MS/MS [15]. Our method assessed linearity, LLOQ, intra- and inter-day precision (RSD %), and accuracy (%) across three levels of quality controls, with values calculated and reported in Fig. 2 and Table 1. The linear dependencies of concentration for each lipid standard are shown by the overlap of XIC (Fig. 2). The calibration curves (Fig. 2, inset) for each lipid were linear within the studied ranges (0.25–25 µM), with mean coefficients of determination (n = 3) of 0.95 or higher. Linearity was evaluated by analyzing a series of standard solutions covering the expected concentration range, and the results demonstrated a satisfactory linear relationship between analyte concentration and the corresponding peak area, consistent with previous studies [23].

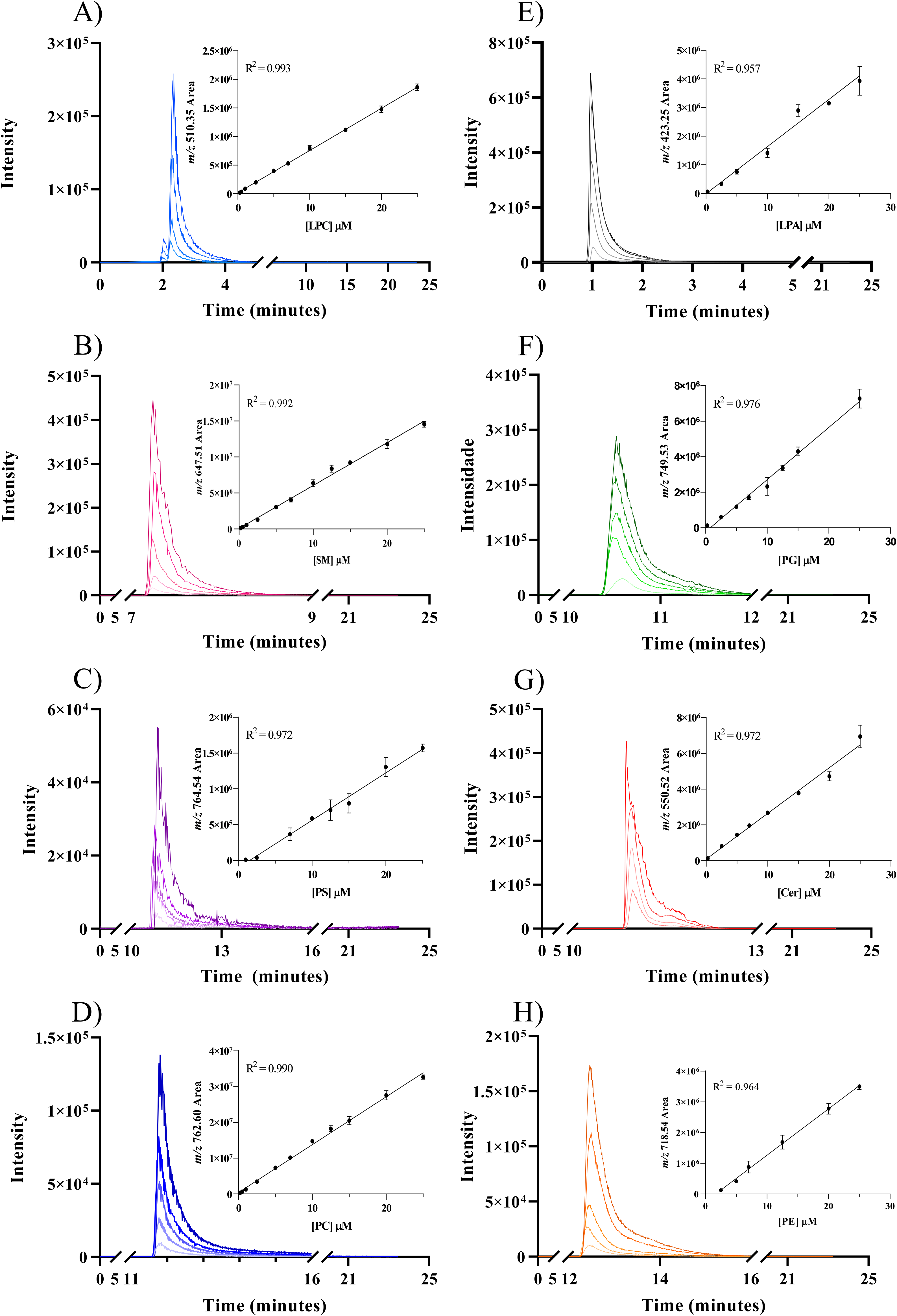

**Table 1.** LLOQ, intra-interday precision and accuracy of the untargeted lipidomic method.

| Lipids species | LLOQ (μM)* | Quality controls* | Intraday (n=5)** |  | Interday (3 days)*** |  |
| --- | --- | --- | --- | --- | --- | --- |
|  |  |  | RSD<sup>1</sup> (%) | Accuracy<sup>2</sup> (%) | RSD<sup>1</sup> (%) | Accuracy<sup>2</sup> (%) |
| LPA (17:1) | 0.25 | LQC | 10.4 | 120 | 10.9 | 115 |
|  |  | MQC | 9.1 | 120 | 13.5 | 119 |
|  |  | HQC | 10.4 | 117 | 9.2 | 116 |
| LPC (17:0) | 0.25 | LQC | 1.40 | 116 | 2.89 | 112 |
|  |  | MQC | 17.0 | 101 | 17.2 | 101 |
|  |  | HQC | 8.62 | 108 | 13.9 | 114 |
| PC (17:0/17:0) | 0.25 | LQC | 0.83 | 86.1 | 1.79 | 85.9 |
|  |  | MQC | 12.3 | 100 | 6.18 | 88.4 |
|  |  | HQC | 5.75 | 85.0 | 4.49 | 81.8 |
| PE (17:0/17:0) | 2.5 | LQC | 13.8 | 112 | 14.9 | 112 |
|  |  | MQC | 2.64 | 118 | 15.3 | 109 |
|  |  | HQC | 17.0 | 101 | 17.2 | 101 |
| PS (17:0/17:0) | 1.0 | LQC | 8.62 | 108 | 13.9 | 114 |
|  |  | MQC | 19.5 | 95.0 | 14.9 | 84.7 |
|  |  | HQC | 14.6 | 96.8 | 11.2 | 85.4 |
| PG (17:0/17:0) | 0.25 | LQC | 10.4 | 120 | 10.9 | 115 |
|  |  | MQC | 9.1 | 120 | 13.5 | 119 |
|  |  | HQC | 10.4 | 117 | 9.2 | 116 |
| SM (18:1/12:0) | 0.25 | LQC | 1.40 | 116 | 2.89 | 112 |
|  |  | MQC | 8.72 | 121 | 7.39 | 118 |
|  |  | HQC | 5.79 | 106 | 4.20 | 105 |
| Cer (18:1/17:0) | 0.25 | LQC | 16.5 | 114 | 15.7 | 114 |
|  |  | MQC | 6.93 | 111 | 5.46 | 116 |
|  |  | HQC | 9.66 | 118 | 8.53 | 114 |

\* Abbreviations: LLOQ, lower limit of quantification; LQC, low-concentration quality control; MQC, Medium-concentration quality control; HQC, high-concentration quality control. \*\* Intra-day precision assessed by analyzing three quality controls at three different concentrations (LQC, MQC and HQC) in quintuplicate. \*\*\* Quality control measurements on three non-consecutive days. <sup>1</sup> Intra-day precision expressed as the relative standard deviation (RSD %) (n=5); <sup>2</sup> Accuracy, determined by back calculation and expressed as coefficient of variation between the average value found by the method and the reference value of the added concentrations.

The LLOQ was defined for each lipid in the mixture (S/N ≥ 10 and RSD ≤ 20%), with values ranging from 0.5 to 2.5 µM. Further, the LLOQ was determined as the lowest concentration that could be measured with acceptable precision and accuracy[15], providing a solid foundation for the quantitative analysis of complex lipid species. The RSD for the replicates was below 20%, the inter lot precision, with RSD (n = 5) of 0.83–19.5 %, was in the range of the accepted criteria, demonstrating the consistency of the method across different sample preparations. The accuracy of the method was evaluated, and the results showed a deviation below 20% of the nominal value, with values ranging from 85 to 120%. This is in accordance with the FDA guidelines which state that the accuracy should be within 20% of the expected value [24,25].

### 3.3. Lipid extraction preparation advances

The efficiency of lipid extraction from biological matrices is a crucial step in the analytical workflow. To address this, we evaluated and optimized the liquid-liquid extraction protocol for application in biological matrices. Internal standard lipid samples were used to validate the extraction protocols. The efficiency of the four lipid extraction protocols (BD, MT, MM, and HD) was assessed by sample recovery (RE) (Table 2).

**Table 2.** Mean recovery of the lipid standards using four extractions protocols in high-concentration quality control level of IS pure samples.

| Average recovery |  |  | BD<sup>a</sup> |  | MT<sup>b</sup> |  | MM<sup>c</sup> |  | HD<sup>d</sup> |  |
| --- | --- | --- | --- | --- | --- | --- | --- | --- | --- | --- |
| Metabolite name | Adduct type | S/N<sup>e</sup> | RE (%)<sup>f</sup> | RSD (%)<sup>g</sup> | RE (%)<sup>f</sup> | RSD (%)<sup>g</sup> | RE (%)<sup>f</sup> | RSD (%)<sup>g</sup> | RE (%)<sup>f</sup> | RSD (%)<sup>g</sup> |
| LPA (17:1) | [M-H]<sup>-</sup> | 1842.7 | 4.63 | 2.24 | 19.48 | 9.67 | 0.85 | 3.19 | 3.39 | 6.43 |
| LPC (17:0) | [M+H]<sup>+</sup> | 500.4 | 51.36 | 4.65 | 48.41 | 13.83 | 36.31 | 18.59 | 49.47 | 9.49 |
| PC (17:0/17:0) | [M+H]<sup>+</sup> | 1098.70 | 52.53 | 7.82 | 94.37 | 7.93 | 93.01 | 14.14 | 52.14 | 9.55 |
| PE (17:0/17:0) | [M-H]<sup>-</sup> | 1395.77 | 82.50 | 13.16 | 61.28 | 9.64 | 28.95 | 13.64 | 29.88 | 0.01 |
| PS (17:0/17:0) | [M+H]<sup>+</sup> | 256.63 | 52.61 | 8.28 | 98.18 | 9.05 | 91.99 | 14.22 | 33.43 | 14.25 |
| PG (17:0/17:0) | [M-H]<sup>-</sup> | 3912.72 | 58.99 | 11.37 | 19.67 | 10.16 | 14.70 | 3.01 | 12.42 | 6.23 |
| SM (18:1/12:0) | [M+H]<sup>+</sup> | 1905.96 | 64.98 | 0.89 | 62.78 | 6.23 | 76.41 | 16.05 | 77.37 | 16.18 |
| Cer (18:1/17:0) | [M-H]<sup>-</sup> | 15614.48 | 92.67 | 6.92 | 14.23 | 0.51 | 18.52 | 11.47 | 75.85 | 2.37 |

<sup>a</sup>BD, chloroform-extraction protocol; <sup>b</sup>MT, MTBE-extraction protocol; <sup>c</sup>MM, MTBE- extraction protocol modified; <sup>d</sup>HD, hybrid-extraction protocol. <sup>e</sup>Relation signal/noise (average, N=5); <sup>f</sup> recovery (RE) was expressed as the percentage of the expected value, calculated by the average area ratio between the extraction samples and the standard samples. <sup>g</sup> relative standard deviation (RSD %) (n=5). A RE value of >100% indicates ionization enhancement, and a value of <100% indicates ionization suppression.

The HD lipid-extraction protocol was newly developed in this work to extract a wide range of analytes with different polarities allowing increasing the coverage of determined lipid species in a sample. This methodology combines chloroform and sequential MTBE extraction, enabling capture of a broader spectrum of lipid species. Additionally, a modification to the MT protocol, known as MM, was introduced, which focuses on increasing the polar fraction and improving the solubilization process [15]. Both methods showed low recoveries for the polar lipid LPA. For LPC, the recovery was around 50% for BD, MT, and HD extraction methods. PC and PS showed the highest recoveries for MT and MM methods, with 94.37% and 93% for PC and 98% and 92% for PS, respectively. PE, PG, and Cer exhibited the expressive values for BD method, with recoveries of 82.5%, 58.99%, and 92.67%, respectively. For SM, the recovery was above 64% using the BD, MT, MM, and HD methods. Thus, it can be concluded that the highest efficiency values for PC, PS, and SM recovery were obtained with the MM methodology. However, for use in complex matrices with a broad distribution of lipid concentrations, the HD extraction protocols may be required to ensure high coverage of both polar and nonpolar lipids in the lipidome.

The HD extraction protocol offers several advantages over other traditional lipid extraction methods, capturing a diverse range of lipid species, including those with varying polarities, thereby enhancing the coverage of determined lipid species in a sample [26]. Furthermore, the MM modification, which targets the polar fraction and improves solubilization, can further expand the range of lipids that can be effectively extracted and analyzed [27].

The implementation of these optimized extraction protocols, coupled with the advancements in mass spectrometry technology, has enabled researchers to conduct comprehensive lipidomic analyses, leading to a deeper understanding of lipid metabolism and its role in various health and disease states [1]. These developments have paved the way for more accurate and efficient lipidomic profiling, ultimately contributing to the advancement of our knowledge in this rapidly evolving field.

Matrix effects were investigated to evaluate the efficiency of the extraction steps and to avoid quantification errors during validation and further analyses. Matrix-dependent recovery was established using plasma samples. The extraction of lipids from complex biological matrices, such as plasma, is a crucial step in lipid analysis, as it can significantly impact the accuracy and reproducibility of the subsequent quantification. Various extraction techniques have been developed to address the matrix complexity and improve the coverage of the lipidome [28]. The values obtained in our experimental approach were given in Table 3.

**Table 3.** Matrix effects of the lipid standards using four extractions protocols in high-concentration quality control level of IS in plasma samples.

| Matrix effects |  |  | BD<sup>a</sup> |  | MT<sup>b</sup> |  | MM<sup>c</sup> |  | HD<sup>d</sup> |  |
| --- | --- | --- | --- | --- | --- | --- | --- | --- | --- | --- |
| Metabolite name | Adduct type | S/N<sup>e</sup> | RE (%)<sup>f</sup> | RSD (%)<sup>g</sup> | RE (%)<sup>f</sup> | RSD (%)<sup>g</sup> | RE (%)<sup>f</sup> | RSD (%)<sup>g</sup> | RE (%) | RSD (%) |
| LPA (17:1) | [M-H]<sup>-</sup> | 1874.35 | 4.01 | 8.39 | 37.83 | 3.20 | 15.11 | 15.24 | 4.42 | 4.82 |
| LPC (17:0) | [M+H]<sup>+</sup> | 1091 | 43.55 | 13.77 | 77.58 | 1.41 | 38.17 | 5.51 | 64.14 | 1.70 |
| PC (17:0/17:0) | [M+H]<sup>+</sup> | 2695.45 | 97.40 | 1.34 | 80.85 | 7.25 | 65.18 | 3.45 | 69.40 | 10.01 |
| PE (17:0/17:0) | [M-H]<sup>-</sup> | 1254.14 | 93.30 | 1.54 | 87.44 | 8.75 | 53.60 | 8.09 | 102.47 | 2.60 |
| PS (17:0/17:0) | [M+H]<sup>+</sup> | 139.45 | 78.39 | 4.18 | 95.17 | 6.18 | 36.32 | 6.10 | 50.36 | 8.37 |
| PG (17:0/17:0) | [M-H]<sup>-</sup> | 3837.69 | 71.68 | 4.52 | 100.09 | 0.61 | 60.65 | 2.27 | 86.37 | 9.16 |
| SM (18:1/12:0) | [M+H]<sup>+</sup> | 2723.20 | 51.42 | 2.02 | 83.81 | 12.60 | 40.32 | 1.46 | 70.04 | 14.31 |
| Cer (18:1/17:0) | [M-H]<sup>-</sup> | 14305.65 | 100.45 | 3.24 | 104.53 | 10.42 | 99.96 | 2.99 | 101.40 | 2.19 |

<sup>a</sup>BD, chloroform-extraction protocol; <sup>b</sup>MT, MTBE-extraction protocol; <sup>c</sup>MM, MTBE- extraction protocol modified; <sup>d</sup>HD, hybrid-extraction protocol. <sup>e</sup>Relation signal/noise (average, N=5); <sup>f</sup> Matrix effect was calculated by the ratio of the mean peak area of an analyte spiked before-extraction to the mean peak area of the same analyte standards multiplied by 100. <sup>g</sup> relative standard deviation (RSD %) (n=5). A RE value of >100% indicates ionization enhancement, and a value of <100% indicates ionization suppression.

The matrix effect results observed for the plasma samples demonstrated an increase in the S/N ratio for the LPC, SM and PC approximately 50 % and decrease for PS. As expected, the matrix effects can affect compounds negatively or positively, lowering or exalting the signal [29]. Recovery of the SM reduced 13.56% in BD method and 36% in MM method compared without matrix. However, improve 21% in the MT method. For PC we observed the similar effects, reduced 14% in MT and 28.8% in MM methods, however in BD to an enhancement of 44.8%. The recovery for PS also increased by 25.8% in the BD method and decreased by 56% in MT method. The RSD deviation was below 15% for all lipids, as recommended by the FDA for matrix effect.

According to the FDA guidelines [19], all data evaluated were acceptable with confidence, precision and accuracy and can be applied for studies of untargeted lipidomics to enable its use in biomarker discovery.

### 3.4 Lipidomics profile of healthy human plasma samples

As proof-of-concept study, the developed and validate method was initially applied to identify and quantify untargeted lipids in healthy human plasma. The four extraction lipids methods were conducted and compared based on the lipidomics annotation. All the lipids species identified for each extraction method carried out were described in Table S3. Many features were obtained in a human plasma sample processed through our workflow. The developed LC-MS/MS method detected 225 molecular lipids between positive and negative modes of analysis (136 for positive, 89 for negative), Table S3 and S4. However, some of these species were not annotated in one specific ionization mode, such as PC, that were detected as protonated molecular species [M+ H]^+^ in the positive ion mode, and as formate adducts [M + HCOO]^−^ in the negative ion mode. As expected, the main classes annotated were PC, LPC and PE (Fig. 3 C and D). The developed and validated method was successful in comprehensively profiling the lipidome of human plasma, providing valuable insights into the diversity of lipids present in the circulatory system [15,30]. Further research is needed to investigate the physiological contributions of these diverse lipids and how their levels change in response to diseases and various therapies.

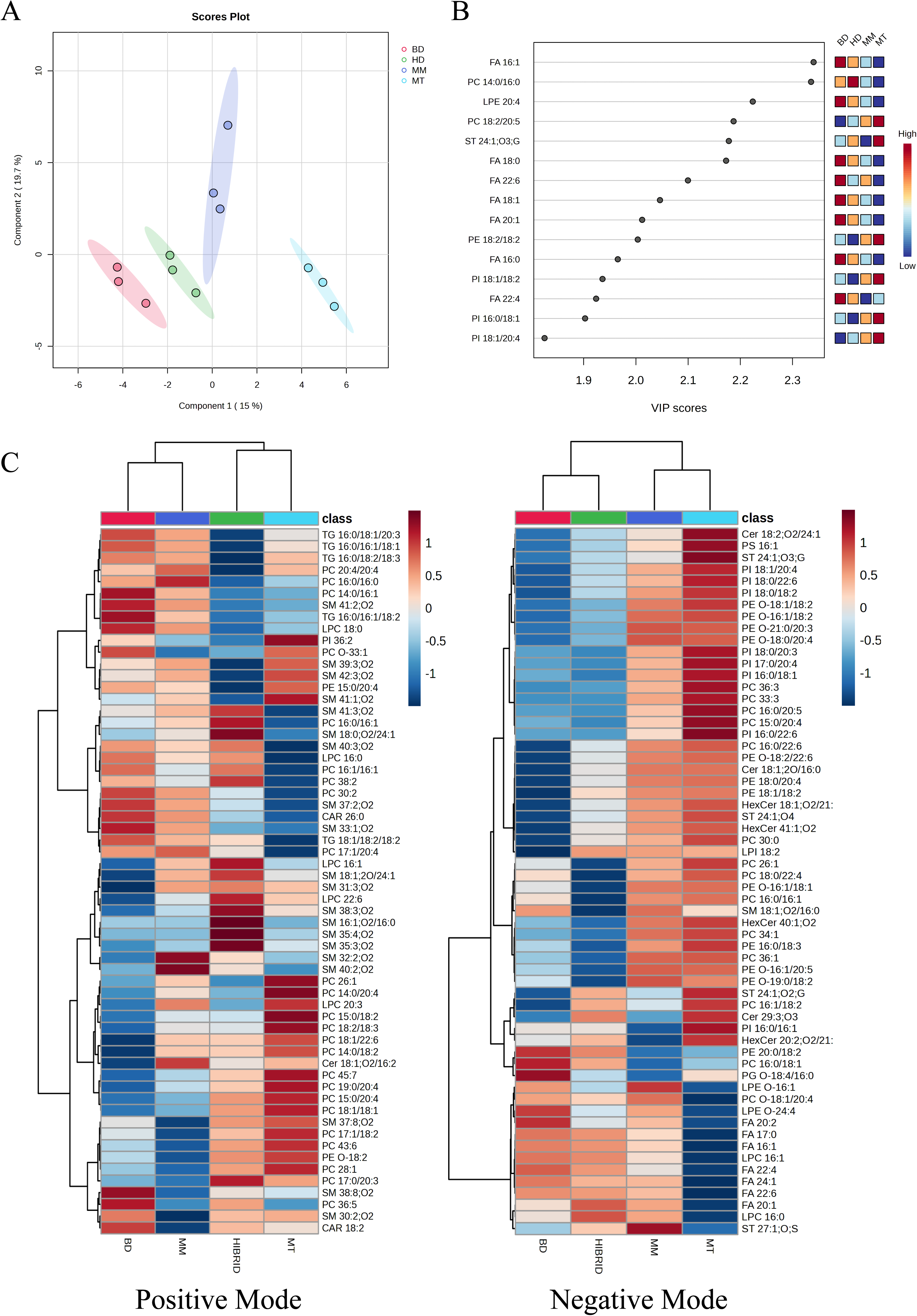

Data obtained by lipids extractions protocols (BD, MT, MM and HD) were compared and the partial least square discriminant analysis (PLS-DA) was applied as an explorative analysis (Fig. 3A). The score plot showed that samples were clustered with respect to their corresponding experimental group, suggesting that each extraction protocol demonstrated a distinct lipidome profile. The variable importance in projection (VIP) score was then utilized to identify the most predictive and differentiating lipid features that could aid in sample classification (Fig. 1B). Lipids are hydrophobic molecules that are soluble in organic solvents but have limited solubility in water. The choice of solvent system is crucial for the efficient extraction of lipids from biological matrices, as it depends on the composition and polarity of the lipids present [31]. The BD method, which employs a chloroform-methanol-water system, has been widely used for the extraction of lipids from a variety of plant and animal tissues [31]. Alternative solvent systems have also been investigated for their efficacy in extracting lipids from sources like black soldier fly [32].The results of the present study demonstrate the importance of selecting an appropriate lipid extraction protocol to obtain a comprehensive representation of the lipidome.

The analysis of the principal lipids species associated with different extraction methods is represented in the heatmap plots allowing for a rapid visual assessment of the similarities and differences between methods. The heatmap shows the abundance of top-60 metabolites in all individuals, for negative (Fig. 3C) and positive mode (Fig. 3D) of LC-MS/MS analysis. The BD and MT extraction methodologies were totally different mainly between TG, DG and PC classes. BD proved to be the best extraction method for TG/DG and MT for PC. Furthermore, MM has an unsatisfactory result for nonpolar molecules, but the HD demonstrates greater capacity to extract SM, also maintaining good performance for PC, LPC and PS, and relatively lower for some PI.

Quantification of lipids identified was carried out and the relative abundance graph (Fig. S1), which demonstrates the extraction efficiency through quantitative values for each class of lipids obtained after extraction. Donut shape graphs showed the relative distribution of individual lipid subclasses in total lipids of healthy individuals’ plasma using BD, MT, MM, and HD extraction protocol. This result provides an excellent overview of the relative abundances in plasma lipidomics by different extraction approaches, demonstrating a similar distribution regardless of the method used, with small variations as expected due to the extraction efficiency studied here. In general, the distribution of total lipids class by relative concentrations in total lipids was similar for both extraction methods evaluated. PC, LPC and PE were the lipid subclasses of highest abundance, accounting for 78.43% to BD, 81.1% to HD, 79.41 to MT, and 79.8% to MM of total lipid concentrations. Other lipids classes quantified of smallest abundance were SM, ST, PS and PI (Fig. S1).

Considering terms of absolute lipid mass, it is possible to state that all protocols had approximately the same mass range between 2000 and 2500 µM, except for the MT method (4284 µM). This difference can be due to the organic phase volume that represents the double compared to that of other methods and consequently more lipids could be extracted by MT method. In contrast, the MM method proved to be the least efficient in absolute values, 2131.94 µM, that also can be explained by the extraction method which has the smallest apolar fraction that was obtained at the end extraction process. BD and HD methods are shown to be very similar, with small variations in absolute mass terms. Based on our results, MT and HD methodologies demonstrated greater diversity of classes and abundance, on the other hand, MM presented a very disadvantaged panel in relation to the other methods.

For the use of extraction methods in untargeted lipidomics, it is necessary that the method cover the widest possible range of classes. Then analyzing all protocols, it is evident that MM and HD demonstrate the greatest diversity of well-extracted lipid classes, with caveats only for some long-chain fatty acids, only MM, and some PC that contain Arachidonic acid (AA) 20:4, Docosahexaenoic acid (DHA) 22:6, and Docosatetraenoic acid (DTA) 22:4. Indeed, we observed that all methodologies of extraction used in this work were able to extract lipids from the main lipid classes known in plasma, however, their differences may be important for less abundant classes, but some biomarkers can be validated.

Thus, there is no single extraction method that is able to extract all lipid classes from biological matrices and careful evaluation and optimization of lipid extraction methods is recommended in order to achieve highest possible extraction efficiencies, throughput and repeatability.

### 3.5 Lipidomics profile of macrophage-derived EVs

From application perspective, the untargeted lipidomics workflow developed and validated was applied to lipidomics profiling in macropagic EVs.

EVs, including exosomes, have emerged as versatile tools for diagnostic and therapeutic medical applications due to their nanoscale sizes, biological origins, various functions, and unique lipid and protein compositions [10,33]. Previous studies have reported the importance of the lipid composition of EVs and its influence on their mechanism of action [34,35]. For instance, changes in the lipidomic profile of EVs have been shown to impact the progression of diseases like ovarian malignancies and prostate cancer [36]. The developed LC-MS/MS methodology enabled detection of a total of one hundred and twenty-four lipid species in positive and negative modes, which can also be included in the 6 subclasses present in EVs macrophage samples. All the anotted lipids species, in positive and negative modes, were demonstrated in Table S5 and S6, respectively.

To EVs samples were performed ultrasonication-treatments previously to the extraction assay to disaggregate EVs after isolation and then evaluate your effects in the lipids species obtained, Fig. S2. The PLS-DA showed the cluster of analytes influenced by the ultrasonic-treatment time before the extraction assay (Fig. 4A and Fig. S2A). Differential lipid distributions were visualized using heatmaps (Fig. S2B) and PLS-DA plots (Fig. S2C) based on their variable importance in projection (VIP) values, which revealed distinct patterns across different ultrasonic treatment durations. This demonstrated an increase in the number of detected lipid species as the treatment time increased, as observed through a sequential membrane trapping (MT) extraction procedure. However, an elevated concentration of free-fatty acids was detected after 60 seconds of ultrasonic treatment, suggesting degradation of complex lipid species. Consequently, treatment duration of 25 seconds was selected for further experimentation. Importantly, the total lipid species recovered from EVs was significantly impacted by the 25-second ultrasonic treatment, likely due to the disruption of lipid and protein aggregation, which can potentially sequester lipids in the aqueous phase during the extraction process.

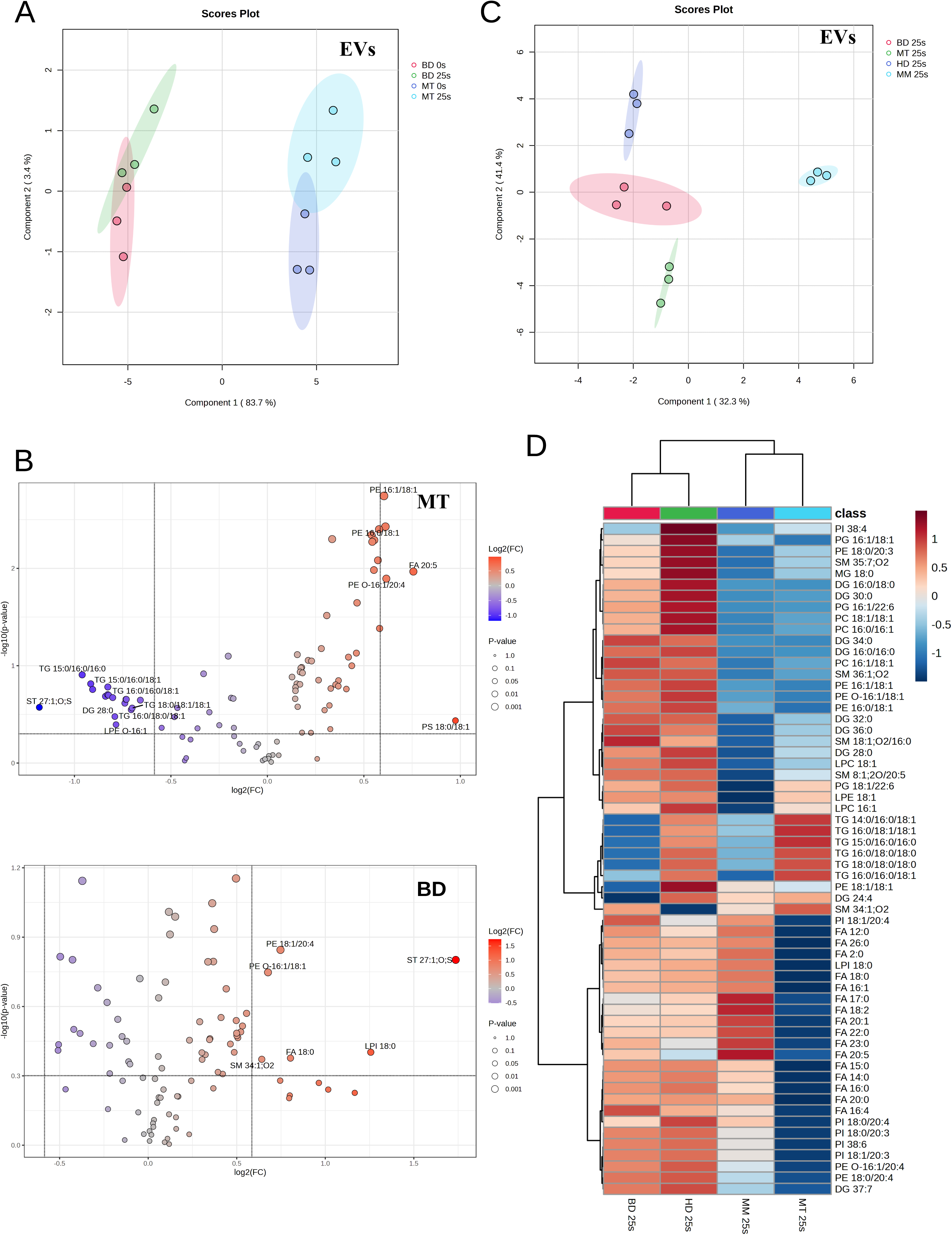

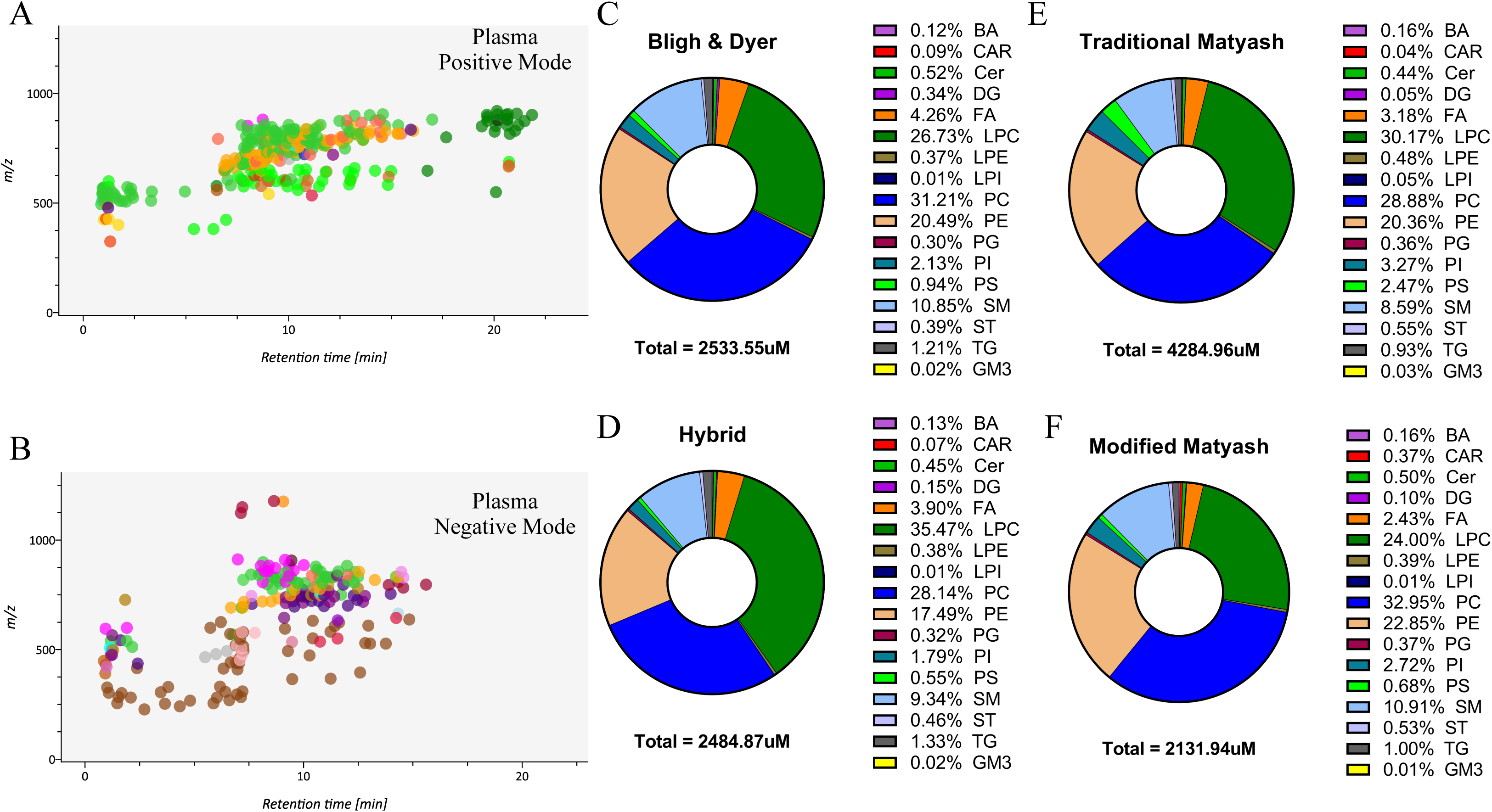

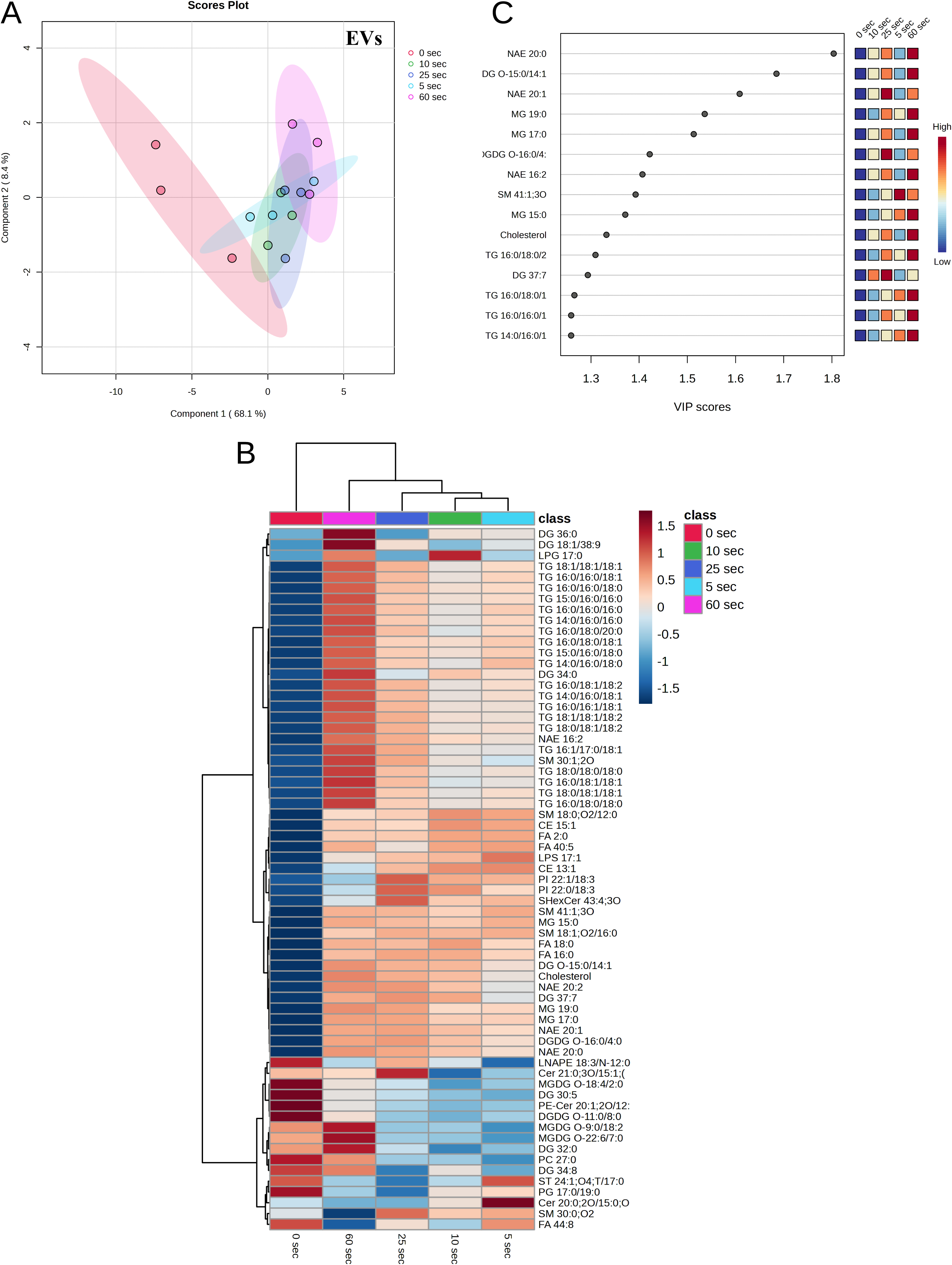

Hierarchical clustering analysis showed an increase in the intensity of lipid species across positive and negative ionization modes following 25 seconds of ultrasonic treatment and various extraction methods, Fig. 4D. Notably, BD protocols exhibited excellent efficiency, except for DG species and PC species in the HD method (Fig. 4D). The HD extraction technique demonstrated superior efficiency in recovering N-acylethanolamines (NAEs), phosphatidylglycerols (PGs), and DG, among other lipid classes. In contrast, the MM protocol’s increased aqueous fraction yielded an intermediate level compared to the BD and MT extremes. The MT method showed low efficiency in detecting PI and FAs, good efficiency for LPEs, and intermediate efficiency for other classes. Overall, the HD extraction appears to be the most suitable approach for comprehensive untargeted lipidomic analysis of extracellular vesicle samples, as it provides efficient recovery of a diverse range of lipid species.

Although the LC-MS/MS approach is used in lipid analysis, there are still limitations related to reproducibility and extraction efficiency that need to be optimized. Variation in identification of lipid species between different methods represents a potential problem. To evaluate the most reliable lipid profile, in this study, we performed a lipid analysis based on untargeted LC-MS/MS for EVs matrix, comparing workflows and evaluating their efficiency in detecting and quantifying lipids present in extracellular vesicles. Our analysis successfully identified 124 lipid species from lipid classes in EVs. It is challenging to compare our results with other reports, since the lipid composition of EVs can be affected by cell type, growth conditions, isolation methods, and lipid analysis [12]. Comprehensive lipid analyzes of EVs demonstrate that the elevated enrichment of cholesterol, SM, glycosphingolipids such as HexCer and LacCer and the decrease in PC are different from their cellular origins [37,38].

Previous studies have suggested that the degree of fatty acid saturation in exosomes and parent cells influences the enrichment of phospholipid species, primarily due to increased levels of PC 14:0/16:0 and 16:0/16:0 [37]. Additionally, PG and PA were detected in low amounts, implying that PA may play a role in EV formation [38]. The high concentrations of CE and TG in EVs could indicate the presence of contaminants, such as lipid droplets and complexed lipoproteins, during EV isolation. The heterogeneity of EVs and their coexistence in cell culture media contribute to the observed differences in lipid composition [39]. Furthermore, low-power sonication can affect EVs, leading to the analysis of lipoprotein-like structures in ultrasonicated EVs that exhibit bilayer lipid droplet-like structures [40,41]. In the other hand, significant changes in the lipid content of EVs were described after treatment with polyunsaturated fatty acids such as arachidonic acid (20:4), eicosapentoic acid (EPA; 20:5) and docosahexaenoic acid (DHA; 22:6), demonstrating that the membrane of EVs could be modulated, favoring the production of pro-resolution lipid mediators [42].

## 4. Conclusions

The untargeted lipidomic workflow herein reported offers an innovative methodology that combines pre-analytical lipids extractions techniques with high-resolution LC-MS/MS to enhance the accuracy and reproducibility of lipidomics research. Lipidomics now plays a pivotal role in biomarker discovery, early disease detection, and the understanding of lipid-mediated signaling pathways. The detailed classification of lipids highlights their structural diversity and functional significance. Traditional lipid extraction techniques have evolved, with more sophisticated methods addressing previous limitations and enhancing recovery and quantitative accuracy. Overall, the advancements in lipidomics, particularly in pre-analytical extraction and mass spectrometry techniques, have significantly enhanced the field’s potential to contribute to personalized medicine and therapeutic development. As the technology and methodologies continue to evolve, lipidomics is poised to uncover even more about the complex roles of lipids in human health and disease.

## CRediT authorship contribution statement

**Adriana Ferreira Lopes Vilela:** Writing – original draft, methodology, investigation, formal analysis, conceptualization. **Miguel Rocha Patricio:** Laboratorial procedure, development of extraction method, data analysis, figure and graph design, table assemblement. **Thiago Felippe:** Table assemblement, graph and figure design. **Viviani Nardini Takahashi:** Methodology. **Nathan Hiroshi Pontes Nakao:** Data analysis, figure and graph design **Jonatan Carvalho:** Laboratorial procedure and data analysis. **Pedro Nobre:** Methodology, investigation, data analysis. **Daniel Lopes:** Data analysis. **Bianca T. M. Oliveira:** Methodology. **Pedro Vieira da Silva-Neto:** Conceptualization, Visualization, Writing – original draft. **Ana Paula Morais Fernandes:** Funding acquisition, Resoucers. **FaustoAlmeida:** Methodology, Resources. **Lucia Helena Faccioli:** Supervision, project administration. **Carlos Arterio Sorgi:** Writing – review & editing, supervision, project administration, funding acquisition, conceptualization.

## Supporting information

Untarget lipidomics_Supplementary information

## Acknowledgements

This work was supported by the São Paulo Research Foundation (FAPESP) [grant numbers 2021/04590-3], National Council for Technological and Scientific Development (CNPq) and Coordinating Development of Higher Level (CAPES). Additional support was given by the National Council for Scientific and Technological Development (CNPq) [grant number 314358/2021-8]. Center of Excellence for Quantification and Identification of Lipids – (CEQIL). A.F.L.V., M.R.P. and C.A.S. also acknowledge CNPq for scholarships [grant numbers 2022/152943 and 2023/2469].

