## Supplementary material for "Optimizing Lipidomics Analysis Workflows for Biological Fluids and Extracellular Vesicles with Integrated Liquid Chromatography Tandem Mass Spectrometry Approaches": Untarget lipidomics_Supplementary information

**Table S1**

List of synthetic lipid standards.

| Common name | Company |
|---|---|
| 1-heptadecanoyl-glycero-3-phosphate (LPA) 17:0 | Avanti Polar Lipids #857127P |
| 1-heptadecanoyl- 2-hydroxy-sn-glycero-3-phosphocholine (LPC) 17:0 | Avanti Polar Lipids #855676P |
| 1,2-diheptadecanoyl-sn-glycero-3-phosphocholine (PC) 17:0/17:0 | Avanti Polar Lipids #850360P |
| 1,2-diheptadecanoyl-sn-glycero-3-phosphoethanolamine (PE) 17:0/17:0 | Avanti Polar Lipids #830756P |
| 1,2-diheptadecanoyl-sn-glycero-3-phospho-1-glycerol (PG) 17:0/17:0 | Avanti Polar Lipids #830456P |
| 1,2-diheptadecanoyl-sn-glycero-3-phospho-L-serine (PS) 17:0/17:0 | Avanti Polar Lipids #840028P |
| N-lauroyl-D-erythro-sphingosylphosphorycholine (SM) 12:0 | Avanti Polar Lipids #860583P |
| N-heptadecanoyl-D-erythro-sphingosine (Cer) 17:0 | Avanti Polar Lipids #860517P |

**Supplementary Table 2**. Peak area at 25 µM, S/N ratio and mass spectrometry parameters for lipids standards.

| **Standards** | **Positive mode** | | | **Negative mode** | | |
|---|---|---|---|---|---|---|
| | Area [25 µM]* | S/N** | $R^{2***}$ | Area [25 µM]* | S/N** | $R^{2***}$ |
| LPA | ND | ND | ND | | 9176.6 ± 9 | 0.9578 |
| LPC | 1.84E+07 | 2498.2 ± 4 | 0.9930 | ND | ND | ND |
| PC | 3.27E+07 | 8326.7 ± 17 | 0.990 | 6.56E+06 | 3423.8 ± 4 | 0.9676 |
| PE | 2.57E+06 | 785.67 ± 25 | 0.986 | 4.05E+06 | 4169.5 ± 46 | 0.8865 |
| PS | 1.57E+06 | 383.8 ± 10 | 0.972 | 4.39E+06 | 3085.5 ± 34 | 0.9233 |
| PG | 4.63E+05 | 77.1 ± 3 | 0.979 | 6.69E+06 | 8896.6 ± 6 | 0.9537 |
| SM | 1.46E+07 | 8731.7 ± 12 | 0.992 | 3.86E+06 | 6632.1 ± 12 | 0.9458 |
| Cer | 2.62E+06 | 98.7 ± 8 | 0.962 | 6.40E+06 | 34039.1 ± 7 | 0.9301 |

*Peak area obtained of each reference standard at 25 µM; **Relation signal/noise; *** linearity of calibration curve. ND, not determined.

**Supplementary Table 3.** Number of lipids identified for robust quantitation in human plasma samples in positive ionization mode.

| Sample Label | Adduct type | BD[a] | BD[b] | BD[c] | MT[a] | MT[b] | MT[c] | MM[a] | MM[b] | MM[c] | HD[a] | HD[b] | HD[c] |
|---|---|---|---|---|---|---|---|---|---|---|---|---|---|
| CAR 18:2 | [M+H]+ | 0.069 | 0.094 | 0.072 | 0.036 | 0.039 | 0.043 | 0.066 | 0.069 | 0.048 | 0.038 | 0.050 | 0.060 |
| CAR 26:0 | [M+H]+ | 0.375 | 0.670 | 0.427 | 0.133 | 0.141 | 0.218 | 2.007 | 1.923 | 0.899 | 0.421 | 0.026 | 0.029 |
| Cer d18:1/16:2 | [M+H]+ | 0.585 | 1.027 | 0.630 | 0.183 | 0.164 | 0.251 | 0.441 | 0.374 | 0.290 | 0.250 | 0.293 | 0.348 |
| Cer d18:1/17:0 | [M+H]+ | 0.008 | 0.027 | 0.010 | 0.002 | 0.003 | 0.006 | 0.007 | 0.004 | 0.004 | 0.004 | 0.003 | 0.002 |
| DGDG O-18:2 | [M+Na]+ | 0.971 | 2.424 | 1.090 | 0.181 | 0.187 | 0.176 | 0.489 | 0.291 | 0.294 | 0.296 | 0.370 | 0.549 |
| LPC 16:0 | [M+H]+ | 44.297 | 33.342 | 35.973 | 26.723 | 27.676 | 24.131 | 22.392 | 20.494 | 0.468 | 34.768 | 34.122 | 35.447 |
| LPC 16:1 | [M+H]+ | 1.265 | 1.065 | 1.014 | 0.482 | 0.494 | 0.476 | 0.238 | 0.205 | 12.217 | 0.707 | 0.846 | 0.868 |
| LPC 17:0 | [M+H]+ | 1.000 | 1.000 | 1.000 | 1.000 | 1.000 | 1.000 | 1.000 | 1.000 | 1.000 | 1.000 | 1.000 | 1.000 |
| LPC 17:2 | [M+H]+ | 2.288 | 1.663 | 1.859 | 1.406 | 1.468 | 1.252 | 1.277 | 1.237 | 0.224 | 1.810 | 1.769 | 1.849 |
| LPC 18:0 | [M+H]+ | 23.699 | 18.148 | 18.755 | 14.335 | 14.341 | 12.726 | 12.956 | 12.766 | 1.473 | 18.099 | 18.480 | 18.903 |
| LPC 20:3 | [M+H]+ | 1.844 | 1.667 | 1.510 | 0.701 | 0.741 | 0.682 | 0.862 | 0.864 | 1.403 | 1.003 | 1.226 | 1.199 |
| LPC 20:4 | [M+Na]+ | 0.663 | 0.582 | 0.579 | 0.657 | 0.718 | 0.618 | 0.220 | 0.239 | 0.318 | 0.630 | 0.448 | 0.483 |
| LPC 20:4 | [M+H]+ | 5.517 | 4.646 | 4.434 | 2.054 | 2.064 | 1.877 | 2.286 | 2.372 | 0.547 | 2.948 | 3.500 | 3.562 |
| LPC 22:6 | [M+H]+ | 1.075 | 0.977 | 1.001 | 0.460 | 0.475 | 0.440 | 0.520 | 0.518 | 2.137 | 0.598 | 0.740 | 0.731 |
| PC 26:1 | [M+H]+ | 0.194 | 0.135 | 0.171 | 0.355 | 0.368 | 0.325 | 0.201 | 0.213 | 0.309 | 0.390 | 0.367 | 0.259 |
| PC 26:1 | [M+Na]+ | 0.027 | 0.020 | 0.028 | 0.052 | 0.056 | 0.050 | 0.033 | 0.032 | 0.045 | 0.057 | 0.059 | 0.041 |
| PC 28:1 | [M+H]+ | 0.416 | 0.276 | 0.351 | 0.777 | 0.810 | 0.739 | 0.434 | 0.547 | 0.575 | 0.599 | 0.606 | 0.526 |
| PC 14:0/16:0 | [M+H]+ | 0.354 | 0.328 | 0.302 | 0.017 | 0.023 | 0.500 | 0.410 | 0.391 | 0.373 | 0.359 | 0.411 | 0.371 |
| PC 14:0/16:1 | [M+H]+ | 0.090 | 0.078 | 0.091 | 0.130 | 0.198 | 0.142 | 0.100 | 0.131 | 0.136 | 0.139 | 0.154 | 0.106 |
| PC 30:2 | [M+H]+ | 0.711 | 0.553 | 0.651 | 1.311 | 1.406 | 1.349 | 0.803 | 0.927 | 0.985 | 1.033 | 1.133 | 0.788 |
| PC 16:0/16:0 | [M+H]+ | 1.467 | 1.334 | 1.301 | 1.269 | 1.386 | 1.338 | 0.962 | 1.050 | 1.221 | 1.365 | 1.552 | 1.423 |
| PC 16:0/16:1 | [M+H]+ | 0.699 | 0.557 | 0.613 | 1.203 | 1.227 | 1.322 | 0.778 | 0.861 | 0.857 | 0.854 | 0.923 | 0.664 |
| PC 14:0/18:2 | [M+H]+ | 0.003 | 0.185 | 0.197 | 0.366 | 0.350 | 0.374 | 0.243 | 0.277 | 0.263 | 0.250 | 0.295 | 0.225 |
| PC 16:1/16:1 | [M+H]+ | 0.624 | 0.487 | 0.568 | 0.011 | 0.009 | 1.263 | 0.714 | 0.854 | 0.403 | 0.024 | 0.993 | 0.662 |
| PC 15:0/18:2 | [M+H]+ | 0.521 | 0.434 | 0.465 | 0.772 | 0.839 | 0.771 | 0.557 | 0.595 | 0.581 | 0.569 | 0.689 | 0.466 |
| PC 17:0/17:0 | [M+H]+ | 1.000 | 1.000 | 1.000 | 1.000 | 1.000 | 1.000 | 1.000 | 1.000 | 1.000 | 1.000 | 1.000 | 1.000 |
| PC 14:0/20:4 | [M+H]+ | 0.348 | 0.295 | 0.336 | 0.712 | 0.793 | 0.766 | 0.413 | 0.484 | 0.507 | 0.526 | 0.023 | 0.386 |
| PC 17:1/18:2 | [M+H]+ | 0.189 | 0.182 | 0.187 | 0.319 | 0.326 | 0.306 | 0.218 | 0.243 | 0.236 | 0.230 | 0.286 | 0.177 |
| PC 15:0/20:4 | [M+H]+ | 0.183 | 0.157 | 0.168 | 0.318 | 0.342 | 0.369 | 0.209 | 0.246 | 0.248 | 0.250 | 0.278 | 0.187 |
| PC 18:1/18:1 | [M+H]+ | 17.861 | 15.843 | 14.703 | 36.191 | 36.469 | 42.854 | 19.412 | 20.410 | 26.341 | 31.324 | 25.959 | 18.041 |
| PC 36:5 | [M+Na]+ | 0.082 | 0.086 | 0.084 | 0.117 | 0.119 | 0.132 | 0.112 | 0.119 | 0.105 | 0.092 | 0.103 | 0.089 |
| PC 18:2/18:3 | [M+H]+ | 1.584 | 1.359 | 1.345 | 2.794 | 2.926 | 3.002 | 1.789 | 2.070 | 2.097 | 2.119 | 2.259 | 1.546 |

| | | | | | | | | | | | | | |
|---|---|---|---|---|---|---|---|---|---|---|---|---|---|
| PC 14:0/22:6 | [M+H]+ | 0.117 | 0.087 | 0.107 | 0.260 | 0.276 | 0.266 | 0.137 | 0.166 | 0.181 | 0.194 | 0.209 | 0.131 |
| PC 17:0/20:3 | [M+H]+ | 0.198 | 0.206 | 0.199 | 0.234 | 0.262 | 0.248 | 0.156 | 0.177 | 0.230 | 0.275 | 0.340 | 0.232 |
| PC 17:0/20:4 | [M+H]+ | 0.672 | 0.676 | 0.615 | 0.772 | 1.214 | 1.180 | 0.797 | 0.855 | 0.884 | 0.909 | 1.010 | 0.717 |
| PC 17:1/20:4 | [M+H]+ | 0.634 | 0.619 | 0.607 | 1.138 | 1.198 | 1.203 | 0.693 | 0.792 | 0.857 | 0.912 | 0.996 | 0.684 |
| PC 38:2 | [M+H]+ | 0.914 | 0.851 | 0.875 | 1.053 | 1.136 | 1.063 | 1.057 | 1.149 | 0.963 | 0.806 | 0.916 | 0.749 |
| PC 16:0/22:4 | [M+H]+ | 10.041 | 8.587 | 8.436 | 13.176 | 13.941 | 14.665 | 7.636 | 8.749 | 10.700 | 12.339 | 12.759 | 9.201 |
| PC 38:6 | [M+Na]+ | 0.109 | 0.141 | 0.117 | 0.137 | 0.141 | 0.161 | 0.145 | 0.150 | 0.120 | 0.096 | 0.122 | 0.102 |
| PC 19:0/20:4 | [M+H]+ | 0.151 | 0.141 | 0.144 | 0.291 | 0.301 | 0.289 | 0.169 | 0.189 | 0.221 | 0.247 | 0.257 | 0.175 |
| PC 18:1/22:6 | [M+H]+ | 0.522 | 0.471 | 0.489 | 0.889 | 0.902 | 0.854 | 0.601 | 0.627 | 0.607 | 0.590 | 0.718 | 0.504 |
| PC 20:4/20:4 | [M+H]+ | 0.192 | 0.171 | 0.180 | 0.374 | 0.419 | 0.422 | 0.223 | 0.261 | 0.269 | 0.276 | 0.315 | 0.210 |
| PC 21:0/20:4 | [M+H]+ | 0.085 | 0.063 | 0.082 | 0.159 | 0.171 | 0.149 | 0.086 | 0.103 | 0.127 | 0.148 | 0.144 | 0.097 |
| PC 20:4/22:4 | [M+H]+ | 0.060 | 0.052 | 0.047 | 0.082 | 0.076 | 0.087 | 0.064 | 0.061 | 0.059 | 0.057 | 0.063 | 0.048 |
| PC 43:6 | [M+H]+ | 0.166 | 0.149 | 0.160 | 0.321 | 0.350 | 0.335 | 0.181 | 0.208 | 0.248 | 0.282 | 0.289 | 0.194 |
| PC 45:7 | [M+H]+ | 0.041 | 0.034 | 0.037 | 0.072 | 0.081 | 0.074 | 0.042 | 0.046 | 0.056 | 0.064 | 0.067 | 0.042 |
| PC O-33:1 | [M+H]+ | 0.688 | 0.549 | 0.499 | 0.518 | 0.534 | 0.563 | 0.511 | 0.585 | 0.272 | 0.009 | 0.383 | 0.419 |
| PC O-36:6 | [M+H]+ | 0.133 | 0.126 | 0.129 | 0.139 | 0.143 | 0.136 | 0.133 | 0.142 | 0.139 | 0.136 | 0.043 | 0.131 |
| PE 15:0/20:4 | [M+H]+ | 0.087 | 0.085 | 0.082 | 0.098 | 0.094 | 0.098 | 0.063 | 0.082 | 0.085 | 0.087 | 0.102 | 0.079 |
| PE O-18:2 | [M+H]+ | 1.917 | 1.447 | 1.706 | 3.395 | 3.855 | 3.773 | 1.697 | 1.867 | 2.571 | 3.163 | 3.213 | 2.384 |
| PI 36:2 | [M+NH4]+ | 0.017 | 0.023 | 0.018 | 0.027 | 0.018 | 0.022 | 0.021 | 0.028 | 0.021 | 0.015 | 0.017 | 0.015 |
| PS 16:1 | [M+H]+ | 0.311 | 0.266 | 0.397 | 1.025 | 0.964 | 1.174 | 1.048 | 1.051 | 1.151 | 1.220 | 1.003 | 1.077 |
| PS 17:0/17:0 | [M+H]+ | 1.000 | 1.000 | 1.000 | 1.000 | 1.000 | 1.000 | 1.000 | 1.000 | 1.000 | 1.000 | 1.000 | 1.000 |
| SM 30:1 | [M+Na]+ | 1.000 | 1.000 | 1.000 | 1.000 | 1.000 | 1.000 | 1.000 | 1.000 | 1.000 | 1.000 | 1.000 | 1.000 |
| SM d18:12/12:0 | [M+H]+ | 1.000 | 1.000 | 1.000 | 1.000 | 1.000 | 1.000 | 1.000 | 1.000 | 1.000 | 1.000 | 1.000 | 1.000 |
| SM 30:2 | [M+H]+ | 0.047 | 0.054 | 0.050 | 0.045 | 0.042 | 0.048 | 0.043 | 0.048 | 0.041 | 0.038 | 0.044 | 0.048 |
| SM 31:3 | [M+H]+ | 107.135 | 5.051 | 5.697 | 4.902 | 4.770 | 4.808 | 5.333 | 5.289 | 6.022 | 6.438 | 5.812 | 6.266 |
| SM d8:0/24:0 | [M+H]+ | 5.308 | 5.804 | 5.791 | 5.615 | 5.535 | 5.260 | 5.577 | 4.762 | 6.632 | 8.123 | 7.773 | 5.951 |
| SM d16:1/16:0 | [M+H]+ | 0.442 | 0.466 | 0.494 | 0.466 | 0.470 | 0.433 | 0.461 | 0.398 | 0.549 | 0.668 | 0.640 | 0.495 |
| SM 32:2 | [M+H]+ | 1.089 | 1.070 | 1.214 | 1.293 | 1.442 | 1.245 | 1.027 | 0.984 | 1.468 | 1.908 | 1.799 | 1.305 |
| SM 33:1 | [M+H]+ | 0.905 | 0.764 | 0.975 | 1.385 | 1.436 | 1.269 | 0.928 | 0.904 | 1.557 | 2.353 | 1.872 | 1.422 |
| SM 35:2 | [M+H]+ | 0.365 | 0.328 | 0.465 | 0.997 | 1.027 | 0.880 | 0.564 | 0.563 | 1.045 | 1.738 | 1.409 | 0.910 |
| SM 35:3 | [M+H]+ | 1.133 | 1.059 | 1.242 | 1.414 | 1.405 | 1.150 | 1.080 | 1.125 | 1.598 | 1.989 | 1.838 | 1.584 |
| SM 35:4 | [M+H]+ | 4.872 | 4.808 | 5.644 | 5.619 | 5.248 | 4.911 | 4.599 | 4.617 | 6.321 | 7.638 | 7.159 | 6.346 |
| SM 37:2 | [M+H]+ | 0.685 | 0.508 | 0.658 | 1.169 | 1.221 | 1.110 | 0.582 | 0.623 | 1.218 | 2.196 | 1.803 | 1.219 |
| SM 37:3 | [M+H]+ | 1.204 | 0.989 | 1.303 | 1.802 | 1.859 | 1.457 | 1.396 | 1.450 | 2.122 | 2.710 | 2.278 | 1.909 |
| SM 37:8 | [M+H]+ | 1.747 | 1.553 | 1.807 | 87.045 | 1.699 | 1.495 | 1.609 | 1.529 | 2.150 | 2.654 | 2.280 | 2.185 |
| SM 38:2 | [M+H]+ | 1.335 | 1.026 | 1.336 | 2.320 | 2.428 | 2.157 | 1.224 | 1.301 | 2.486 | 4.325 | 3.581 | 2.489 |

| | | | | | | | | | | | | | |
|---|---|---|---|---|---|---|---|---|---|---|---|---|---|
| SM 38:3 | [M+H]+ | 4.000 | 2.966 | 3.967 | 4.970 | 5.217 | 4.448 | 3.679 | 3.979 | 6.037 | 7.960 | 6.189 | 5.867 |
| SM 38:8 | [M+H]+ | 0.064 | 0.055 | 0.072 | 0.104 | 0.101 | 0.087 | 0.094 | 0.091 | 0.112 | 0.126 | 0.112 | 0.101 |
| SM 39:2 | [M+H]+ | 0.371 | 0.276 | 0.356 | 0.705 | 0.647 | 0.614 | 0.305 | 0.335 | 0.679 | 1.302 | 1.041 | 0.726 |
| SM 39:3 | [M+H]+ | 0.434 | 0.333 | 0.433 | 0.593 | 0.619 | 0.545 | 0.425 | 0.423 | 0.648 | 0.861 | 0.824 | 0.658 |
| SM 40:2 | [M+H]+ | 0.851 | 0.640 | 0.769 | 1.543 | 1.419 | 1.418 | 0.652 | 0.792 | 1.599 | 3.053 | 2.542 | 1.535 |
| SM 40:3 | [M+H]+ | 2.685 | 2.225 | 2.522 | 3.803 | 4.025 | 3.581 | 2.530 | 2.570 | 3.936 | 5.237 | 4.882 | 3.799 |
| SM 41:1 | [M+H]+ | 4.429 | 3.414 | 3.992 | 7.491 | 7.158 | 7.239 | 3.298 | 4.094 | 8.103 | 14.929 | 13.044 | 7.567 |
| SM 41:2 | [M+H]+ | 0.647 | 0.497 | 0.590 | 1.126 | 1.019 | 1.058 | 0.507 | 0.604 | 1.191 | 2.183 | 1.855 | 1.138 |
| SM 41:3 | [M+H]+ | 0.377 | 0.294 | 0.368 | 0.428 | 0.419 | 0.377 | 0.369 | 0.366 | 0.495 | 0.593 | 0.517 | 0.492 |
| SM d18:0/24:1 | [M+H]+ | 2.076 | 1.595 | 1.981 | 3.289 | 3.305 | 3.080 | 1.950 | 1.921 | 3.237 | 4.760 | 4.417 | 3.260 |
| SM d18:1/24:1 | [M+H]+ | 0.290 | 0.215 | 0.274 | 0.469 | 0.451 | 0.417 | 0.264 | 0.269 | 0.460 | 0.691 | 0.589 | 0.481 |
| SM 42:3 | [M+H]+ | 5.128 | 4.345 | 5.201 | 8.198 | 7.829 | 7.285 | 5.183 | 5.030 | 8.319 | 11.978 | 10.829 | 8.822 |
| TG 8:0/9:0/13:0 | [M+Na]+ | 0.984 | 1.214 | 0.944 | 1.270 | 1.244 | 1.259 | 0.639 | 0.688 | 0.901 | 1.006 | 1.177 | 1.241 |
| TG 8:0/8:0/19:0 | [M+Na]+ | 0.350 | 0.376 | 0.636 | 0.176 | 0.281 | 0.290 | 0.233 | 0.212 | 0.176 | 0.158 | 0.221 | 0.353 |
| TG 16:0/16:1/18:1 | [M+Na]+ | 0.312 | 0.548 | 0.396 | 0.251 | 0.249 | 0.253 | 0.372 | 0.365 | 0.252 | 0.197 | 0.210 | 0.244 |
| TG 16:0/16:1/18:2 | [M+Na]+ | 0.212 | 0.350 | 0.281 | 0.175 | 0.186 | 0.206 | 0.255 | 0.242 | 0.184 | 0.156 | 0.163 | 0.184 |
| TG 16:0/18:1/18:2 | [M+Na]+ | 0.671 | 1.156 | 0.982 | 0.535 | 0.554 | 0.562 | 0.786 | 0.736 | 0.614 | 0.554 | 0.536 | 0.645 |
| TG 16:0/18:2/18:3 | [M+Na]+ | 0.200 | 0.356 | 0.310 | 0.180 | 0.185 | 0.188 | 0.238 | 0.224 | 0.196 | 0.182 | 0.182 | 0.226 |
| TG 16:0/18:1/20:3 | [M+Na]+ | 0.241 | 0.446 | 0.399 | 0.210 | 0.232 | 0.225 | 0.297 | 0.304 | 0.259 | 0.237 | 0.239 | 0.262 |
| TG 18:1/18:2/18:2 | [M+Na]+ | 0.300 | 0.534 | 0.473 | 0.254 | 0.244 | 0.275 | 0.333 | 0.351 | 0.287 | 0.256 | 0.285 | 0.312 |
| TG O-16:3/18:3/20:5 | [M+Na]+ | 0.550 | 0.956 | 0.811 | 0.416 | 0.429 | 0.471 | 0.596 | 0.605 | 0.494 | 0.439 | 0.472 | 0.506 |

Abbreviations: BD, chloroform-extraction protocol; MT, MTBE-extraction protocol; MM, MTBE-extraction protocol modified. a, b and c: ratio peak area analyte/ peak area internal standard in triplicate. Value = 1 means that are IS samples.

**Supplementary Table 4.** Number of lipids identified for robust quantitation in human plasma samples in negative ionization mode.

| Metabolite name<br>Label | Adduct type | BDa | BDb | BDc | HDa | HDb | HDc | MMa | MMb | MMc | MTa | MTb | MTc |
|---|---|---|---|---|---|---|---|---|---|---|---|---|---|
| BA 24:1;O3;T/17:0 | [M-H]- | 0.2860 | 0.1848 | 0.2482 | 0.3707 | 0.4081 | 0.2852 | 0.4294 | 0.4913 | 1.8750 | 0.6649 | 0.6757 | 0.5997 |
| Cer 29:3;O3 | [M-H]- | 0.0237 | 0.0152 | 0.0164 | 0.0291 | 0.0424 | 0.0208 | 0.0055 | 0.0076 | 0.8750 | 0.0616 | 0.0450 | 0.0392 |
| Cer d18:1/16:0 | [M-H]- | 0.0471 | 0.0453 | 0.0428 | 0.0382 | 0.0490 | 0.0541 | 0.0696 | 0.0733 | 3.2500 | 0.0718 | 0.0680 | 0.0788 |
| Cer d18:0/16:2 | [M-H]- | 0.0149 | 0.0174 | 0.0177 | 0.0143 | 0.0245 | 0.0273 | 0.0253 | 0.0258 | 0.1250 | 0.0249 | 0.0250 | 0.0231 |
| Cer d18:1/17:0 | [M-H]- | 1.0000 | 1.0000 | 1.0000 | 1.0000 | 1.0000 | 1.0000 | 1.0000 | 1.0000 | 1.0000 | 1.0000 | 1.0000 | 1.0000 |
| Cer 36:2;O3 | [M-H]- | 0.0461 | 0.0326 | 0.0338 | 0.0657 | 0.0975 | 0.0460 | 0.0362 | 0.0472 | 5.2500 | 0.1530 | 0.1037 | 0.0877 |

| | | | | | | | | | | | | | |
|---|---|---|---|---|---|---|---|---|---|---|---|---|---|
| Cer d18:2/24:1 | [M-H]- | 0.0704 | 0.0563 | 0.0596 | 0.0890 | 0.0992 | 0.0501 | 0.0731 | 0.0866 | 0.1250 | 0.1078 | 0.1332 | 0.1223 |
| FA 16:0 | [M-H]- | 77.8664 | 50.1747 | 87.0463 | 66.1135 | 83.1958 | 22.1340 | 28.2434 | 37.9505 | 6.2632 | 7.7602 | 8.1098 | 5.8069 |
| FA 16:1 | [M-H]- | 9.4117 | 3.4087 | 8.6961 | 7.2316 | 10.8715 | 1.6422 | 3.5577 | 5.9361 | 0.3355 | 0.9199 | 0.8848 | 0.5239 |
| FA 17:0 | [M-H]- | 3.6228 | 1.8371 | 3.7888 | 2.9831 | 3.9799 | 0.8721 | 1.7994 | 2.2842 | 0.4718 | 0.3806 | 0.4684 | 0.2954 |
| FA 18:0 | [M-H]- | 50.0491 | 30.7113 | 53.2236 | 41.1956 | 46.3266 | 13.4185 | 22.4871 | 33.7148 | 9.9877 | 5.3022 | 6.4642 | 4.4465 |
| FA 18:1 | [M-H]- | 111.0131 | 63.1661 | 109.2393 | 84.0946 | 111.6064 | 27.0685 | 43.7888 | 65.7275 | 0.7556 | 15.1011 | 14.3030 | 10.3623 |
| FA 20:1 | [M-H]- | 1.6049 | 0.8109 | 1.4977 | 1.1185 | 1.4844 | 0.3906 | 0.7199 | 1.0981 | 0.0463 | 0.1915 | 0.2335 | 0.1491 |
| FA 20:2 | [M-H]- | 2.1203 | 1.0478 | 1.9960 | 1.5623 | 2.2701 | 0.4065 | 0.8322 | 1.2495 | 0.0036 | 0.2314 | 0.2561 | 0.1724 |
| FA 20:4 | [M-H]- | 3.9682 | 2.3698 | 4.0419 | 3.0966 | 3.4579 | 0.9557 | 2.2647 | 3.3360 | 0.0020 | 0.6340 | 0.6405 | 0.4793 |
| FA 22:4 | [M-H]- | 2.3812 | 1.4713 | 2.3458 | 1.7261 | 2.5402 | 0.5438 | 0.9308 | 1.3297 | 0.0151 | 0.3296 | 0.3151 | 0.2291 |
| FA 22:6 | [M-H]- | 2.3775 | 1.1920 | 2.1127 | 1.6128 | 2.9263 | 0.6617 | 1.5182 | 1.9287 | 0.6491 | 0.2104 | 0.4383 | 0.2108 |
| FA 24:1 | [M-H]- | 0.5412 | 0.3260 | 0.4976 | 0.2844 | 0.4159 | 0.2072 | 0.3219 | 0.4681 | 0.0019 | 0.0709 | 0.0795 | 0.0574 |
| FA 27:0 | [M-H]- | 4.1568 | 2.2804 | 3.8788 | 2.9574 | 3.7445 | 0.9535 | 0.6506 | 0.8120 | 0.0068 | 0.5202 | 0.5035 | 0.3624 |
| FA 30:5 | [M-H]- | 2.5163 | 1.4246 | 2.5074 | 1.8772 | 2.3635 | 0.5552 | 0.2931 | 0.2706 | 0.0163 | 0.3157 | 0.2979 | 0.2153 |
| FA 32:8 | [M-H]- | 1.1235 | 0.5744 | 0.8723 | 0.9337 | 1.1108 | 0.2926 | 0.1027 | 0.1532 | 0.0256 | 0.1597 | 0.1524 | 0.0913 |
| FA 34:6 | [M-H]- | 3.2013 | 1.8085 | 3.0754 | 3.7982 | 4.3451 | 1.0661 | 0.3025 | 0.4315 | 0.0012 | 0.8154 | 0.7304 | 0.4805 |
| FA 36:3 | [M-H]- | 0.1321 | 0.1051 | 0.1378 | 0.0982 | 0.1049 | 0.0560 | 0.1258 | 0.1254 | 0.0312 | 0.0163 | 0.0182 | 0.0184 |
| FA 36:4 | [M-H]- | 0.3482 | 0.3603 | 0.4242 | 0.2555 | 0.2351 | 0.2360 | 0.4100 | 0.3485 | 0.5214 | 0.0413 | 0.0492 | 0.0490 |
| FA 36:7 | [M-H]- | 4.3167 | 2.2670 | 4.2081 | 5.0268 | 5.5521 | 1.3183 | 0.4542 | 0.6212 | 0.0146 | 1.1211 | 0.9899 | 0.6806 |
| FA 36:8 | [M-H]- | 3.7170 | 1.9611 | 3.4113 | 4.2154 | 5.0361 | 1.1982 | 0.5020 | 0.5924 | 0.0072 | 1.0410 | 0.8629 | 0.5939 |
| FA 38:8 | [M-H]- | 2.1078 | 1.1166 | 1.9260 | 2.0388 | 2.5241 | 0.6065 | 0.5781 | 0.7981 | 0.0135 | 0.4269 | 0.3835 | 0.2749 |
| FA 40:5 | [M-H]- | 0.9603 | 1.0410 | 1.6600 | 1.0048 | 1.3039 | 0.5791 | 1.3604 | 1.4254 | 0.0654 | 0.0938 | 0.1973 | 0.1854 |
| FA 40:9 | [M-H]- | 12.2409 | 6.0397 | 11.0527 | 12.2843 | 14.9982 | 3.2686 | 5.5234 | 8.6124 | 0.0008 | 2.5219 | 2.1899 | 1.5341 |
| FA 42:10 | [M-H]- | 1.5885 | 0.9726 | 1.8815 | 1.3525 | 1.9142 | 0.6072 | 0.8080 | 1.0244 | 0.1323 | 0.2131 | 0.2510 | 0.1592 |
| FA 42:12 | [M-H]- | 0.6239 | 0.3596 | 0.6408 | 0.5815 | 0.6022 | 0.2339 | 0.3709 | 0.4305 | 0.0034 | 0.1128 | 0.1140 | 0.0794 |
| FA 42:5 | [M-H]- | 0.9413 | 0.6993 | 0.9931 | 1.2349 | 1.1423 | 0.4366 | 1.6911 | 1.7894 | 0.0797 | 0.1040 | 0.2759 | 0.1989 |
| FA 42:6 | [M-H]- | 1.1773 | 0.8243 | 0.9049 | 0.5004 | 0.4668 | 0.2828 | 0.7061 | 0.7132 | 0.0150 | 0.1268 | 0.1155 | 0.0716 |
| FA 44:11 | [M-H]- | 0.9980 | 0.5276 | 0.9282 | 0.7009 | 1.0204 | 0.3155 | 0.4135 | 0.5346 | 0.0040 | 0.1517 | 0.1693 | 0.1083 |
| FA 44:12 | [M-H]- | 0.8540 | 0.7355 | 0.7291 | 0.2552 | 0.3128 | 0.2659 | 0.5222 | 0.5006 | 0.0276 | 0.0866 | 0.0764 | 0.0436 |
| FA 44:8 | [M-H]- | 2.9045 | 2.2347 | 2.1412 | 0.7999 | 0.7858 | 0.5231 | 0.9784 | 1.0006 | 0.0150 | 0.2388 | 0.1990 | 0.1098 |
| GM3 32:1;O2 | [M-H]- | 0.0286 | 0.0285 | 0.0289 | 0.0287 | 0.0347 | 0.0194 | 0.0142 | 0.0199 | 7.5000 | 0.0557 | 0.0598 | 0.0523 |
| GM3 34:2;O2 | [M-H]- | 0.0778 | 0.0637 | 0.0691 | 0.0556 | 0.0822 | 0.0486 | 0.0353 | 0.0434 | 16.5000 | 0.0902 | 0.0944 | 0.1081 |

| | | | | | | | | | | | | | |
|---|---|---|---|---|---|---|---|---|---|---|---|---|---|
| Hex2Cer 34:1;O2 | [M+HCOO]- | 0.0208 | 0.0221 | 0.0223 | 0.0253 | 0.0274 | 0.0255 | 0.0388 | 0.0383 | 3.2500 | 0.0371 | 0.0351 | 0.0331 |
| HexCer d18:1/16:0 | [M+HCOO]- | 0.3970 | 0.3827 | 0.3664 | 0.3835 | 0.4931 | 0.5739 | 0.5357 | 0.6059 | 7.5000 | 0.6903 | 0.6567 | 0.6526 |
| HexCer d18:1/21:1;O | [M+HCOO]- | 0.0462 | 0.0310 | 0.0373 | 0.0505 | 0.0507 | 0.0318 | 0.0580 | 0.0651 | 3.2500 | 0.0975 | 0.0957 | 0.0644 |
| HexCer 40:1;O2 | [M-H]- | 0.0732 | 0.0482 | 0.0595 | 0.0791 | 0.0814 | 0.0542 | 0.0881 | 0.1032 | 0.7500 | 0.1709 | 0.1600 | 0.1089 |
| HexCer 41:1;O2 | [M-H]- | 0.0325 | 0.0223 | 0.0262 | 0.0317 | 0.0339 | 0.0265 | 0.0362 | 0.0423 | 1.6250 | 0.0674 | 0.0570 | 0.0384 |
| HexCer d20:2/21:1;O | [M+HCOO]- | 0.0489 | 0.0399 | 0.0456 | 0.0713 | 0.0831 | 0.0396 | 0.0635 | 0.0783 | 4.6250 | 0.1070 | 0.1137 | 0.0873 |
| HexCer 42:2;O2 | [M-H]- | 0.0766 | 0.0571 | 0.0678 | 0.1033 | 0.1173 | 0.0635 | 0.0970 | 0.1132 | 2.1250 | 0.1531 | 0.1725 | 0.1355 |
| LPA 17:1 | [M-H]- | 1.0000 | 1.0000 | 1.0000 | 1.0000 | 1.0000 | 1.0000 | 1.0000 | 1.0000 | 1.0000 | 1.0000 | 1.0000 | 1.0000 |
| LPC 16:0 | [M+HCOO]- | 62.7900 | 31.8095 | 62.4595 | 70.4959 | 68.0204 | 19.4767 | 35.7455 | 47.4408 | 0.0001 | 12.9368 | 12.8963 | 9.9135 |
| LPC 16:1 | [M+HCOO]- | 2.1186 | 0.9202 | 1.7692 | 1.8737 | 1.8860 | 0.4917 | 0.9151 | 1.3135 | 0.0003 | 0.3405 | 0.3556 | 0.2726 |
| LPE 18:2 | [M-H]- | 13.5443 | 6.5979 | 12.8576 | 12.3337 | 13.3334 | 3.5865 | 9.7362 | 13.4760 | 0.0145 | 2.7562 | 2.8118 | 2.1959 |
| LPE 20:4 | [M-H]- | 10.0699 | 5.4189 | 9.5595 | 8.0882 | 9.3421 | 2.4558 | 6.7670 | 9.8972 | 0.0005 | 1.8181 | 1.9060 | 1.4911 |
| LPE O-16:1 | [M-H]- | 1.0153 | 0.4814 | 0.9350 | 1.3520 | 1.3673 | 0.3477 | 0.9761 | 1.3451 | 0.0080 | 0.2874 | 0.3277 | 0.2254 |
| LPE O-24:4 | [M-H]- | 3.9259 | 2.2812 | 4.1378 | 4.6220 | 4.3629 | 1.2595 | 2.3476 | 3.2566 | 0.0003 | 0.8471 | 0.8361 | 0.6234 |
| LPI 18:0 | [M-H]- | 0.5118 | 0.4627 | 0.7602 | 0.5684 | 0.4503 | 0.1592 | 0.4160 | 0.5474 | 0.0030 | 0.3425 | 0.4464 | 0.2483 |
| LPI 18:2 | [M-H]- | 0.4079 | 0.1804 | 0.3115 | 0.3537 | 0.3234 | 0.1005 | 0.2501 | 0.2890 | 0.0516 | 0.2766 | 0.2927 | 0.1882 |
| PC 26:1 | [M+HCOO]- | 0.3234 | 0.2622 | 0.3118 | 0.4326 | 0.4584 | 0.2481 | 0.3069 | 0.3925 | 1.3636 | 0.4753 | 0.4896 | 0.4576 |
| PC 30:0 | [M+HCOO]- | 3.7364 | 2.8476 | 3.4165 | 3.6239 | 4.3786 | 2.6012 | 3.9153 | 4.4792 | 0.9091 | 5.4448 | 6.4185 | 6.1284 |
| PC 16:0/16:1 | [M+HCOO]- | 0.4846 | 0.4683 | 0.4625 | 0.3908 | 0.4878 | 0.4934 | 0.5989 | 0.6074 | 1.7273 | 0.6854 | 0.6484 | 0.6646 |
| PC 14:0/18:2 | [M+HCOO]- | 0.6223 | 0.5166 | 0.5666 | 0.5684 | 0.7055 | 0.4773 | 0.6554 | 0.8113 | 0.2727 | 0.8877 | 0.9783 | 1.0118 |
| PC 33:3 | [M+HCOO]- | 0.6015 | 0.5705 | 0.5799 | 0.5364 | 0.6274 | 0.5977 | 0.7256 | 0.7739 | 0.0455 | 0.9033 | 0.8958 | 0.9305 |
| PC 17:0/17:0 | [M+HCOO]- | 1.0000 | 1.0000 | 1.0000 | 1.0000 | 1.0000 | 1.0000 | 1.0000 | 1.0000 | 1.0000 | 1.0000 | 1.0000 | 1.0000 |
| PC 34:1 | [M+HCOO]- | 2.7703 | 2.9154 | 2.6656 | 2.1366 | 2.7033 | 3.0283 | 3.5486 | 3.5154 | 11.1818 | 3.6398 | 3.5360 | 3.8636 |
| PC 16:0/18:1 | [M+HCOO]- | 9.3076 | 9.9962 | 9.2757 | 7.5643 | 9.4021 | 10.0413 | 12.0785 | 12.3025 | 1.5909 | 13.1241 | 12.7320 | 14.0373 |
| PC 16:0/18:2 | [M+HCOO]- | 29.7454 | 32.6775 | 29.5802 | 22.5475 | 28.8068 | 34.9177 | 39.6644 | 39.7833 | 9.3182 | 40.1939 | 37.8125 | 41.4794 |
| PC 16:1/18:2 | [M+HCOO]- | 1.8202 | 1.5046 | 1.6420 | 1.6694 | 2.0066 | 1.4364 | 1.9000 | 2.2138 | 0.5455 | 2.6121 | 2.7703 | 2.8931 |
| PC 14:0/20:4 | [M+HCOO]- | 0.2968 | 0.2351 | 0.2655 | 0.2902 | 0.3445 | 0.2073 | 0.2826 | 0.3589 | 0.2727 | 0.4328 | 0.4591 | 0.4764 |
| PC 15:0/20:4 | [M+HCOO]- | 0.2397 | 0.2185 | 0.2229 | 0.2015 | 0.2509 | 0.1903 | 0.2481 | 0.2808 | 0.4091 | 0.3338 | 0.3526 | 0.3831 |
| PC 36:1 | [M+HCOO]- | 0.1166 | 0.0942 | 0.1359 | 0.0972 | 0.1269 | 0.0789 | 0.1508 | 0.1587 | 0.0455 | 0.1068 | 0.1781 | 0.1724 |
| PC 18:0/18:1 | [M+HCOO]- | 2.4775 | 2.5710 | 2.4141 | 2.1244 | 2.5919 | 0.0349 | 0.0468 | 3.1083 | 1.1818 | 3.3652 | 3.3926 | 0.0555 |
| PC 18:0/18:2 | [M+HCOO]- | 16.1724 | 16.9122 | 15.8188 | 13.2855 | 16.4998 | 17.2408 | 19.5454 | 20.3265 | 8.8182 | 21.7711 | 21.8110 | 23.3649 |

| | | | | | | | | | | | | | |
|---|---|---|---|---|---|---|---|---|---|---|---|---|---|
| PC 36:3 | [M+HCOO]- | 0.2494 | 0.2338 | 0.2457 | 0.2369 | 0.2799 | 0.2333 | 0.2937 | 0.3177 | 0.7727 | 0.3565 | 0.3764 | 0.3931 |
| PC 16:0/20:4 | [M+HCOO]- | 11.5329 | 11.3246 | 10.9043 | 8.4356 | 11.0350 | 12.1672 | 13.6061 | 14.2450 | 10.6818 | 15.1905 | 14.4284 | 15.7136 |
| PC 18:2/18:2 | [M+HCOO]- | 3.2631 | 0.1259 | 2.8679 | 2.7977 | 3.4195 | 2.5103 | 3.2553 | 3.7989 | 48.2273 | 4.3584 | 4.6329 | 4.9091 |
| PC 16:0/20:5 | [M+HCOO]- | 2.2103 | 1.8715 | 2.0266 | 1.9479 | 2.3689 | 1.6972 | 2.2676 | 2.6627 | 1.9091 | 3.1450 | 3.2726 | 3.3933 |
| PC 38:3 | [M+HCOO]- | 1.2846 | 1.2672 | 1.1665 | 1.0427 | 1.2700 | 1.2683 | 1.5156 | 1.5471 | 0.5909 | 1.6992 | 1.6782 | 1.7778 |
| PC 18:0/20:3 | [M+HCOO]- | 3.5880 | 3.4829 | 3.4266 | 3.1197 | 3.7229 | 3.3039 | 4.0321 | 4.1507 | 1.5455 | 4.7399 | 4.9754 | 5.2498 |
| PC 16:0/22:6 | [M+HCOO]- | 4.4448 | 3.6203 | 3.9967 | 4.1054 | 4.8518 | 3.7421 | 4.4659 | 4.8785 | 3.5455 | 6.5283 | 6.6020 | 6.8741 |
| PC 18:2/20:5 | [M+HCOO]- | 0.3000 | 0.2305 | 0.2630 | 0.2911 | 0.3500 | 0.2119 | 0.2908 | 0.3501 | 0.1364 | 0.4069 | 0.4646 | 0.4849 |
| PC 18:0/22:4 | [M+HCOO]- | 0.5239 | 0.4932 | 0.4993 | 0.4572 | 0.5542 | 0.4519 | 0.5574 | 0.5891 | 0.6364 | 0.7041 | 0.7514 | 0.7769 |
| PC 18:0/22:6 | [M+HCOO]- | 1.5663 | 1.5581 | 1.5178 | 1.3768 | 1.6339 | 1.6140 | 1.8533 | 1.9233 | 0.7727 | 2.0968 | 2.2585 | 2.3889 |
| PC 20:4/20:4 | [M+HCOO]- | 0.2582 | 0.2140 | 0.2335 | 0.2388 | 0.3011 | 0.1879 | 0.2570 | 0.3017 | 0.4091 | 0.3614 | 0.3839 | 0.3870 |
| PC 20:4/22:6 | [M+HCOO]- | 0.0549 | 0.0446 | 0.0497 | 0.0600 | 0.0661 | 0.0371 | 0.0507 | 0.0657 | 0.4091 | 0.0785 | 0.0986 | 0.0950 |
| PC O-18:0/20:4 | [M+HCOO]- | 0.8194 | 0.7378 | 0.7594 | 0.7323 | 0.8846 | 0.7021 | 0.8946 | 0.9306 | 0.4545 | 1.0939 | 1.1810 | 1.2248 |
| PC O-18:1/20:4 | [M+HCOO]- | 0.9852 | 0.9218 | 0.9060 | 0.8279 | 0.4245 | 0.9227 | 1.1591 | 1.1646 | 1.0455 | 1.3577 | 0.1029 | 0.1099 |
| PE 17:0/17:0 | [M-H]- | 1.0000 | 1.0000 | 1.0000 | 1.0000 | 1.0000 | 1.0000 | 1.0000 | 1.0000 | 1.0000 | 1.0000 | 1.0000 | 1.0000 |
| PE 16:0/18:2 | [M-H]- | 0.6655 | 0.9935 | 0.9036 | 0.8640 | 1.2230 | 1.0452 | 1.8469 | 1.8843 | 1.1304 | 1.5438 | 1.9502 | 2.3832 |
| PE 16:0/18:3 | [M-H]- | 0.0619 | 0.1063 | 0.0892 | 0.0825 | 0.1228 | 0.0898 | 0.1489 | 0.1645 | 0.6957 | 0.1414 | 0.2212 | 0.3056 |
| PE 18:0/18:1 | [M-H]- | 0.5319 | 0.8756 | 0.7030 | 0.5836 | 0.8542 | 0.8711 | 1.5055 | 1.4477 | 0.6957 | 1.0124 | 1.2822 | 1.6766 |
| PE 18:0/18:2 | [M-H]- | 1.4923 | 2.4128 | 2.0050 | 1.6622 | 2.3916 | 2.7340 | 4.2537 | 3.8533 | 3.4783 | 3.0557 | 3.8581 | 4.9375 |
| PE 18:1/18:2 | [M-H]- | 0.6063 | 0.8544 | 0.7719 | 0.7016 | 1.0723 | 0.8991 | 1.5213 | 1.5428 | 0.8261 | 1.2747 | 1.6303 | 2.0740 |
| PE 18:2/18:2 | [M-H]- | 0.2833 | 0.3901 | 0.3660 | 0.3300 | 0.4804 | 0.3035 | 0.5937 | 0.7048 | 1.2609 | 0.5711 | 0.9642 | 1.1386 |
| PE 20:0/18:2 | [M-H]- | 5.4554 | 9.0091 | 7.2445 | 6.1298 | 8.8480 | 10.3167 | 14.7665 | 13.4875 | 1.0435 | 10.4519 | 11.2531 | 16.0610 |
| PE 18:0/20:4 | [M-H]- | 2.1589 | 3.2257 | 2.8354 | 2.6617 | 3.9372 | 3.6644 | 6.0603 | 5.9557 | 3.1304 | 4.9255 | 6.1221 | 7.6122 |
| PE 22:5/22:5 | [M-H]- | 0.4717 | 0.7156 | 0.6075 | 0.4961 | 0.7178 | 0.8094 | 1.2080 | 1.1399 | 0.6957 | 0.8842 | 0.9951 | 1.3752 |
| PE O-16:1/18:1 | [M-H]- | 0.2666 | 0.3739 | 0.3452 | 0.3450 | 0.4941 | 0.3713 | 0.6869 | 0.7274 | 2.7826 | 0.5870 | 0.7349 | 0.8971 |
| PE O-16:1/18:2 | [M-H]- | 0.5318 | 0.7165 | 0.7342 | 0.7516 | 1.0706 | 0.8797 | 1.7817 | 1.9078 | 1.3478 | 1.5561 | 2.0089 | 2.3849 |
| PE O-18:1/18:2 | [M-H]- | 1.0821 | 1.3592 | 1.3855 | 1.4497 | 1.9678 | 1.2638 | 2.5417 | 2.8088 | 0.7826 | 2.3432 | 3.2061 | 3.6962 |
| PE O-16:1/20:4 | [M-H]- | 2.5337 | 3.5333 | 3.2803 | 3.2361 | 4.6767 | 3.9702 | 7.6884 | 8.2626 | 0.5217 | 6.6196 | 8.4634 | 10.2010 |
| PE O-16:1/20:5 | [M-H]- | 0.2053 | 0.2973 | 0.2655 | 0.2269 | 0.3667 | 0.3593 | 0.5571 | 0.5352 | 0.0435 | 0.4384 | 0.4820 | 0.6475 |
| PE O-20:0/17:0 | [M-H]- | 0.4320 | 0.7224 | 0.5910 | 0.5586 | 0.6444 | 0.7517 | 1.0033 | 0.8822 | 0.0435 | 0.6236 | 0.6841 | 0.8996 |
| PE O-19:0/18:2 | [M-H]- | 12.1686 | 21.4671 | 16.3974 | 13.0414 | 19.3642 | 0.1780 | 34.8597 | 34.2172 | 6.3478 | 24.3382 | 24.9406 | 35.1391 |
| PE O-20:1/18:2 | [M-H]- | 0.1382 | 0.1831 | 0.1644 | 0.1784 | 0.2470 | 0.1797 | 0.3295 | 0.3483 | 0.2609 | 0.2881 | 0.3820 | 0.4374 |

| | | | | | | | | | | | | | |
|---|---|---|---|---|---|---|---|---|---|---|---|---|---|
| PE O-18:0/20:4 | [M-H]- | 0.5420 | 0.8621 | 0.6795 | 0.5988 | 0.8993 | 0.9830 | 1.3668 | 1.3598 | 0.2609 | 1.0437 | 1.1991 | 1.7049 |
| PE O-18:2/22:6 | [M-H]- | 0.6993 | 0.8708 | 0.8814 | 0.8321 | 1.1973 | 0.9504 | 1.6409 | 1.8198 | 1.0870 | 1.4563 | 1.9383 | 2.2994 |
| PE O-21:0/20:3 | [M-H]- | 0.9428 | 1.5149 | 1.2243 | 1.0883 | 1.5032 | 1.5880 | 2.4700 | 2.1956 | 0.6087 | 1.7736 | 2.0204 | 2.9216 |
| PE-Cer 12:2;2O/16:4 | [M-H]- | 0.8657 | 1.0035 | 0.9702 | 0.9956 | 1.4306 | 0.9684 | 1.2387 | 1.3588 | 0.6087 | 2.3621 | 1.9195 | 2.6171 |
| PG 17:0/17:0 | [M-H]- | 1.0000 | 1.0000 | 1.0000 | 1.0000 | 1.0000 | 1.0000 | 1.0000 | 1.0000 | 1.0000 | 1.0000 | 1.0000 | 1.0000 |
| PG 18:0/18:0;O | [M-H]- | 0.1697 | 0.3039 | 0.1740 | 0.1384 | 0.1114 | 0.2812 | 0.1899 | 0.1073 | 0.1273 | 0.1359 | 0.1120 | 0.1050 |
| PG O-18:4/16:0 | [M-H]- | 0.1243 | 0.1412 | 0.1005 | 0.0785 | 0.1249 | 0.0834 | 0.0282 | 0.0161 | 5.2909 | 0.0502 | 0.0614 | 0.0564 |
| PG O-22:6/16:0 | [M-H]- | 0.0408 | 0.0759 | 0.0481 | 0.0229 | 0.0281 | 0.0650 | 0.0262 | 0.0235 | 0.2182 | 0.0197 | 0.0176 | 0.0279 |
| PI 16:0/16:1 | [M-H]- | 0.1472 | 0.0828 | 0.1091 | 0.1323 | 0.1458 | 0.0714 | 0.1405 | 0.1886 | 0.3636 | 0.2765 | 0.3302 | 0.2874 |
| PI 16:0/18:1 | [M-H]- | 0.5349 | 0.4215 | 0.5402 | 0.4268 | 0.5204 | 0.3538 | 0.7050 | 0.8265 | 0.5455 | 1.1958 | 1.2880 | 1.1017 |
| PI 16:0/18:2 | [M-H]- | 1.0071 | 0.6124 | 0.7520 | 0.7630 | 0.8485 | 0.5045 | 1.0376 | 1.2614 | 1.0000 | 1.7697 | 2.2266 | 1.8915 |
| PI 18:0/18:2 | [M-H]- | 1.1978 | 0.7743 | 1.0720 | 0.9602 | 1.1564 | 0.7448 | 1.4807 | 1.7230 | 1.4091 | 2.5358 | 2.7506 | 2.3308 |
| PI 18:1/18:2 | [M-H]- | 0.1880 | 0.1156 | 0.1502 | 0.1700 | 0.1997 | 0.0890 | 0.1806 | 0.2468 | 0.4545 | 0.3706 | 0.4244 | 0.3691 |
| PI 16:0/20:4 | [M-H]- | 1.5238 | 0.8489 | 1.1092 | 1.1418 | 1.3478 | 0.6750 | 1.3645 | 1.6794 | 0.0455 | 2.3546 | 3.0039 | 2.6204 |
| PI 17:0/20:4 | [M-H]- | 0.0839 | 0.0590 | 0.0689 | 0.0667 | 0.0808 | 0.0528 | 0.0956 | 0.1113 | 0.4545 | 0.1456 | 0.1745 | 0.1534 |
| PI 18:0/20:3 | [M-H]- | 0.7499 | 0.5448 | 0.7169 | 0.6446 | 0.7635 | 0.4645 | 0.9268 | 1.1725 | 2.7273 | 1.5793 | 1.7331 | 1.5219 |
| PI 18:0/20:4 | [M-H]- | 0.0359 | 3.8417 | 5.1412 | 0.0269 | 5.1073 | 3.2767 | 0.0408 | 8.1629 | 0.6818 | 11.0296 | 0.0677 | 10.5634 |
| PI 18:1/20:4 | [M-H]- | 0.6498 | 0.4036 | 0.4843 | 0.4758 | 0.5555 | 0.3148 | 0.5812 | 0.7275 | 0.5455 | 1.0094 | 1.2508 | 1.0966 |
| PI 16:0/22:6 | [M-H]- | 0.1942 | 0.1124 | 0.1502 | 0.1763 | 0.1872 | 0.0935 | 0.1681 | 0.2180 | 0.1818 | 0.3253 | 0.4346 | 0.3491 |
| PI 18:0/22:5 | [M-H]- | 0.0850 | 0.0746 | 0.0672 | 0.0557 | 0.0682 | 0.0630 | 0.1231 | 0.1246 | 326.5909 | 0.0707 | 0.0553 | 0.0546 |
| PI 18:0/22:6 | [M-H]- | 0.2911 | 0.2076 | 0.2760 | 0.3004 | 0.3243 | 0.1573 | 0.3106 | 0.4303 | 1.2727 | 0.5804 | 0.9653 | 0.6757 |
| PS 16:1 | [M-H]- | 0.3712 | 0.6063 | 1.6768 | 0.4257 | 0.9307 | 0.5931 | 0.6919 | 0.9072 | 0.3500 | 2.1925 | 2.6671 | 3.3585 |
| PS 17:0/17:0 | [M-H]- | 1.0000 | 1.0000 | 1.0000 | 1.0000 | 1.0000 | 1.0000 | 1.0000 | 1.0000 | 1.0000 | 1.0000 | 1.0000 | 1.0000 |
| SM d18:0/12:0 | [M+HCOO]- | 0.0883 | 0.0957 | 0.0926 | 0.0831 | 0.0877 | 0.0867 | 0.0870 | 0.0825 | 0.6154 | 0.0835 | 0.0854 | 0.0849 |
| SM d18:1/12:0 | [M+HCOO]- | 1.0000 | 1.0000 | 1.0000 | 1.0000 | 1.0000 | 1.0000 | 1.0000 | 1.0000 | 1.0000 | 1.0000 | 1.0000 | 1.0000 |
| SM d16:1/16:0 | [M+HCOO]- | 1.1856 | 1.4259 | 1.3206 | 1.2687 | 1.2835 | 1.5922 | 1.4850 | 1.4482 | 1.2308 | 1.3035 | 1.5306 | 1.4995 |
| SM d18:1/14:0 | [M+HCOO]- | 1.4550 | 1.7801 | 1.6297 | 1.4792 | 1.2327 | 1.9879 | 1.8386 | 1.8051 | 0.1538 | 1.5843 | 1.8137 | 1.8071 |
| SM 32:2 | [M+HCOO]- | 0.5430 | 0.7826 | 0.6796 | 0.4976 | 0.5272 | 0.8535 | 0.8227 | 0.7135 | 1.1923 | 0.5593 | 0.6755 | 0.6823 |
| SM d14:0/18:2 | [M+HCOO]- | 0.1385 | 0.1363 | 0.1318 | 0.1170 | 0.1284 | 0.1281 | 0.1330 | 0.1375 | 1.1538 | 0.1516 | 0.1569 | 0.1653 |
| SM d18:1/16:0 | [M+HCOO]- | 2.8191 | 4.6718 | 3.6176 | 2.3252 | 2.5226 | 5.7444 | 4.8302 | 3.7880 | 3.5769 | 2.9655 | 2.9158 | 3.1617 |
| SM d19:0/17:1 | [M+HCOO]- | 0.6616 | 1.1358 | 0.7929 | 0.4434 | 0.4632 | 1.0766 | 1.1391 | 0.8643 | 0.7308 | 0.6604 | 0.6063 | 0.6797 |
| SM d18:2/18:0 | [M+HCOO]- | 0.3604 | 0.4969 | 0.3914 | 0.2251 | 0.2720 | 0.5143 | 0.4749 | 0.3400 | 97.6538 | 0.3290 | 0.3416 | 0.3709 |

| | | | | | | | | | | | | | |
|---|---|---|---|---|---|---|---|---|---|---|---|---|---|
| SM 38:2 | [M+HCOO]- | 0.4889 | 0.6178 | 0.5152 | 0.3522 | 0.3629 | 0.5135 | 0.6525 | 0.5530 | 0.0769 | 0.5148 | 0.5143 | 0.4753 |
| SM 38:8 | [M+HCOO]- | 4.1586 | 5.7242 | 4.3064 | 2.5988 | 2.9016 | 5.7163 | 5.3806 | 4.3737 | 2.5000 | 3.9260 | 3.7704 | 4.3572 |
| SM d18:1/22:0 | [M+HCOO]- | 2.4426 | 2.9728 | 2.4678 | 1.5208 | 1.5216 | 2.3144 | 3.2054 | 2.6122 | 2.0769 | 2.5688 | 2.4523 | 2.1461 |
| SM d18:2/24:1 | [M+HCOO]- | 2.8091 | 3.4631 | 2.8606 | 1.9335 | 2.0676 | 3.1583 | 3.3350 | 2.7921 | 1.1154 | 2.6372 | 2.7419 | 2.8454 |
| ST 24:1;O2;G | [M-H]- | 0.0195 | 0.0111 | 0.0141 | 0.0272 | 0.0384 | 0.0242 | 0.0187 | 0.0387 | 1.8750 | 0.0671 | 0.0460 | 0.0448 |
| ST 24:1;O3;G | [M-H]- | 0.0137 | 0.0070 | 0.0079 | 0.0259 | 0.0258 | 0.0207 | 0.0135 | 0.0196 | 43.8750 | 0.2925 | 0.1931 | 0.2023 |
| ST 24:1;O3;T/21:2 | [M-H]- | 0.2228 | 0.1975 | 0.2034 | 0.2422 | 0.2971 | 0.2746 | 0.3075 | 0.3309 | 0.8750 | 0.4144 | 0.0201 | 0.4147 |
| ST 24:1;O4 | [M-H]- | 0.0392 | 0.0213 | 0.0227 | 0.0646 | 0.0912 | 0.0642 | 0.0933 | 0.1228 | 24.7500 | 0.2165 | 0.1440 | 0.1494 |
| ST 24:1;O4/12:0 | [M-H]- | 0.3308 | 0.2242 | 0.2484 | 0.4105 | 0.6097 | 0.2743 | 0.2243 | 0.3284 | 6.8750 | 0.8550 | 0.5792 | 0.5058 |
| ST 27:1;O;S | [M-H]- | 0.2920 | 0.2299 | 0.2878 | 0.5215 | 0.5529 | 0.2316 | 0.6753 | 0.7819 | 70.8750 | 0.7692 | 0.9334 | 0.7631 |

Abbreviations: BD, chloroform-extraction protocol; MT, MTBE-extraction protocol; MM, MTBE-extraction protocol modified. a, b and c: ratio peak area analyte/ peak area internal standard in triplicate. Value = 1 means that are IS samples.

**Supplementary Table 5.** Number of lipids identified for robust quantitation in EVs samples in positive ionization mode.

| Metabolite name<br>Label | Adduct type | BD[a] | BD[b] | BD[c] | HD[a] | HD[b] | HD[c] | MT[a] | MT[b] | MT[c] | MM[a] | MM[b] | MM[c] |
|---|---|---|---|---|---|---|---|---|---|---|---|---|---|
| CE 11:0 | [M+Na]+ | 0 | 0 | 0 | 0 | 0.02 | 0.03 | 0 | 0 | 0 | 0 | 0.02 | 0 |
| CE 13:1 | [M+Na]+ | 1.02 | 1.34 | 0.47 | 0 | 1.1 | 0.51 | 0 | 0 | 0.25 | 0 | 0.17 | 0.19 |
| CE 15:1 | [M+Na]+ | 1.15 | 1.92 | 0.87 | 0.49 | 0.45 | 0.85 | 0 | 0 | 0 | 0 | 0 | 0 |
| Cer d18:1/17:0 | [M+H]+ | 1 | 1 | 1 | 1 | 1 | 1 | 1 | 1 | 1 | 1 | 1 | 1 |
| DG 16:0/16:0 | [M+NH4]+ | 0.04 | 0.03 | 0.04 | 0.03 | 0.04 | 0.13 | 0 | 0 | 0 | 0 | 0 | 0 |
| DG 16:0/18:0 | [M+NH4]+ | 0.1 | 0.09 | 0.05 | 0.08 | 0.08 | 0.22 | 0 | 0 | 0 | 0 | 0 | 0 |
| DG 24:4 | [M+Na]+ | 0 | 0 | 0 | 0.11 | 0.09 | 0.1 | 0 | 0 | 0.04 | 0.04 | 0.04 | 0.04 |
| DG 28:0 | [M+Na]+ | 0.06 | 0.05 | 0.06 | 0.2 | 0.2 | 0.12 | 0 | 0 | 0 | 0 | 0 | 0 |
| DG 30:0 | [M+Na]+ | 0.1 | 0.1 | 0.12 | 0.4 | 0.31 | 0.33 | 0 | 0 | 0 | 0 | 0 | 0 |

| | | | | | | | | | | | | | |
|---|---|---|---|---|---|---|---|---|---|---|---|---|---|
| DG 32:0 | [M+Na]+ | 1.07 | 2.6 | 0.65 | 2.81 | 2.97 | 1.77 | 0 | 0 | 0.01 | 0 | 0 | 0.07 |
| DG 32:0 | [M+Na]+ | 2.13 | 2.42 | 1.65 | 2.61 | 2.82 | 5.4 | 0 | 0 | 0 | 0 | 0.02 | 0.01 |
| DG 34:0 | [M+Na]+ | 0 | 0.08 | 0.13 | 0.02 | 0.11 | 0 | 0 | 0 | 0 | 0 | 0 | 0 |
| DG 34:0 | [M+Na]+ | 3.2 | 3.28 | 2.3 | 3.61 | 3.53 | 7.29 | 0 | 0 | 0.15 | 0 | 0.08 | 0.01 |
| DG 36:0 | [M+Na]+ | 0.69 | 0.72 | 0.21 | 0.11 | 0.16 | 1.24 | 0 | 0 | 0.06 | 0 | 0.03 | 0 |
| DG 37:7 | [M+Na]+ | 59.7 | 75.69 | 94.89 | 307.77 | 240.24 | 159.77 | 0 | 0 | 0 | 0 | 0 | 0 |
| LPC 16:1 | [M+H]+ | 0.25 | 0.31 | 0.25 | 0 | 0.65 | 0.36 | 0 | 0.02 | 0.14 | 0 | 0 | 0 |
| LPC 17:0 | [M+H]+ | 1 | 1 | 1 | 1 | 1 | 1 | 1 | 1 | 1 | 1 | 1 | 1 |
| LPC 17:0 | [M+Na]+ | 1 | 1 | 1 | 1 | 1 | 1 | 1 | 1 | 1 | 1 | 1 | 1 |
| LPC 18:0 | [M+H]+ | 0.29 | 0.29 | 0.3 | 0.03 | 0.19 | 0.17 | 0 | 0 | 0.18 | 0 | 0 | 0 |
| LPC 18:1 | [M+H]+ | 0.95 | 1.04 | 1.08 | 0.69 | 0.86 | 0.72 | 0 | 0 | 0.15 | 0 | 0 | 0 |
| MG 18:0 | [M+NH4]+ | 0.01 | 0.02 | 0.05 | 0.03 | 0.09 | 0.08 | 0 | 0 | 0 | 0 | 0.01 | 0 |
| MGDG O-26:6/3:0 | [M+Na]+ | 5.42 | 5.38 | 4.87 | 22.42 | 16 | 14.97 | 0 | 0 | 0.06 | 0.87 | 0.65 | 1.04 |
| NAE 14:0 | [M+H]+ | 0.04 | 0.03 | 0.06 | 0.24 | 0.2 | 0.12 | 0 | 0 | 0 | 0 | 0 | 0 |
| NAE 16:2 | [M+H]+ | 0.01 | 0.01 | 0.02 | 0.07 | 0.06 | 0.05 | 0 | 0 | 0 | 0 | 0 | 0 |
| NAE 18:2 | [M+H]+ | 0 | 0 | 0.01 | 0.02 | 0.03 | 0.01 | 0 | 0 | 0 | 0 | 0 | 0 |
| NAE 20:0 | [M+H]+ | 0.32 | 0.26 | 0.59 | 1.32 | 1.54 | 0.85 | 0 | 0 | 0 | 0 | 0 | 0 |
| NAE 20:1 | [M+H]+ | 0.02 | 0.01 | 0.04 | 0.05 | 0.06 | 0.04 | 0 | 0 | 0 | 0 | 0 | 0 |
| PC 16:0/18:1 | [M+H]+ | 0.34 | 0.29 | 0.66 | 1.71 | 2.33 | 0.7 | 0 | 0 | 0.1 | 0.14 | 0.1 | 0.1 |
| PC 17:0/17:0 | [M+H]+ | 1 | 1 | 1 | 1 | 1 | 1 | 1 | 1 | 1 | 1 | 1 | 1 |
| PC 17:0/17:0 | [M+H]+ | 1 | 1 | 1 | 1 | 1 | 1 | 1 | 1 | 1 | 1 | 1 | 1 |

| | | | | | | | | | | | | | |
|---|---|---|---|---|---|---|---|---|---|---|---|---|---|
| PC 18:1/18:1 | [M+H]+ | 0.06 | 0.02 | 0.1 | 0.17 | 0.22 | 0.11 | 0 | 0 | 0.01 | 0 | 0 | 0 |
| PC 34:0 | [M+Na]+ | 1 | 1 | 1 | 1 | 1 | 1 | 1 | 1 | 1 | 1 | 1 | 1 |
| PI 38:4 | [M+Na]+ | 0.11 | 0.1 | 0.22 | 0.78 | 0.8 | 0.33 | 0 | 0 | 0.04 | 0.06 | 0.04 | 0.04 |
| SM d18:1/12:0 | [M+H]+ | 1 | 1 | 1 | 1 | 1 | 1 | 1 | 1 | 1 | 1 | 1 | 1 |
| SM d18:1/16:0 | [M+H]+ | 0.24 | 0.26 | 0.3 | 0.07 | 0.2 | 0.17 | 0.01 | 0 | 0.15 | 0.02 | 0.02 | 0.09 |
| SM 30:1;2O | [M+Na]+ | 1 | 1 | 1 | 1 | 1 | 1 | 1 | 1 | 1 | 1 | 1 | 1 |
| SM 34:1;O2 | [M+H]+ | 0.14 | 0.15 | 0.13 | 0.07 | 0.09 | 0.1 | 0 | 0 | 0.02 | 0.01 | 0.01 | 0.01 |
| SM 35:7;O2 | [M+H]+ | 0.04 | 0.04 | 0.04 | 0.16 | 0.2 | 0.09 | 0 | 0 | 0 | 0 | 0 | 0 |
| SM 36:1;O2 | [M+H]+ | 0.08 | 0.08 | 0.06 | 0.06 | 0.06 | 0.08 | 0 | 0 | 0.02 | 0.01 | 0.01 | 0.01 |
| TG 14:0/16:0/16:0 | [M+NH4]+ | 0.08 | 0.05 | 0 | 0.03 | 0.01 | 0.08 | 0.11 | 0.3 | 0 | 0.01 | 0.25 | 0 |
| TG 14:0/16:0/18:1 | [M+NH4]+ | 0.09 | 0.03 | 0.03 | 0.05 | 0.04 | 0.09 | 0 | 0.03 | 0 | 0.05 | 0.55 | 0 |
| TG 15:0/16:0/16:0 | [M+NH4]+ | 0.05 | 0 | 0 | 0.02 | 0 | 0.06 | 0 | 0.06 | 0 | 0 | 0.32 | 0 |
| TG 15:0/16:0/18:1 | [M+NH4]+ | 0.03 | 0 | 0 | 0.04 | 0.03 | 0.04 | 0 | 0 | 0.01 | 0 | 0.29 | 0 |
| TG 16:0/16:0/16:0 | [M+NH4]+ | 0.33 | 0.21 | 0.03 | 0.23 | 0.13 | 0.41 | 0.2 | 0.33 | 0.1 | 0 | 1.1 | 0 |
| TG 16:0/16:0/16:0 | [M+NH4]+ | 0.31 | 0.2 | 0.06 | 0.22 | 0.1 | 0.4 | 0.86 | 0.81 | 0.22 | 0 | 0.99 | 0 |
| TG 16:0/16:0/18:0 | [M+NH4]+ | 0.51 | 0.45 | 0.21 | 0.35 | 0.22 | 0.74 | 0.61 | 0 | 0.63 | 0.12 | 1.53 | 0.11 |
| TG 16:0/16:0/18:1 | [M+NH4]+ | 0.09 | 0.08 | 0.04 | 0.16 | 0.08 | 0.16 | 0 | 0 | 0.02 | 0.03 | 0.97 | 0.13 |
| TG 16:0/16:1/18:1 | [M+NH4]+ | 0 | 0 | 0 | 0 | 0 | 0 | 0 | 0 | 0 | 0 | 0 | 0 |
| TG 16:0/18:0/18:0 | [M+NH4]+ | 0.29 | 0.33 | 0.2 | 0.33 | 0.18 | 0.66 | 0 | 0.3 | 0 | 0 | 1.64 | 0.35 |
| TG 16:0/18:0/18:0 | [M+Na]+ | 0.05 | 0.06 | 0 | 0 | 0.01 | 0.09 | 0 | 0 | 0 | 0 | 0.09 | 0.01 |
| TG 16:0/18:0/18:1 | [M+NH4]+ | 0.08 | 0.01 | 0.04 | 0.07 | 0.04 | 0.17 | 0.03 | 0 | 0 | 0 | 0.57 | 0.06 |

| | | | | | | | | | | | | | |
|---|---|---|---|---|---|---|---|---|---|---|---|---|---|
| TG 16:0/18:1/18:1 | [M+NH4]+ | 0 | 0 | 0 | 0 | 0 | 0.13 | 0 | 0 | 0 | 0 | 0.73 | 0 |
| TG 18:0/18:0/18:0 | [M+NH4]+ | 0.03 | 0.11 | 0 | 0.13 | 0.11 | 0.25 | 0 | 0 | 0 | 0 | 0.39 | 0.15 |
| TG 18:0/18:1/18:1 | [M+NH4]+ | 0 | 0 | 0 | 0 | 0.05 | 0.16 | 0 | 0 | 0 | 0 | 0.76 | 0 |
| TG 8:0/8:0/22:4 | [M+Na]+ | 9.8 | 7.74 | 6.98 | 22.71 | 14.05 | 15.24 | 0 | 0 | 0.05 | 0.87 | 1.18 | 1.31 |
| TG 8:0/8:0/24:4 | [M+Na]+ | 1.28 | 1.87 | 1.33 | 0 | 0.49 | 1.11 | 0 | 0 | 0 | 0 | 0 | 0 |

Abbreviations: BD, chloroform-extraction protocol; MT, MTBE-extraction protocol; MM, MTBE-extraction protocol modified. a, b and c: ratio peak area analyte/ peak area internal standard in triplicate. All these experiments were carried out with 25 seconds of sonication of samples before the extraction protocols. Value = 1 means that are IS samples.

**Supplementary Table 6.** Number of lipids identified for robust quantitation in EVs samples in negative ionization mode.

| **Metabolite name** / **Label** | **Adduct type** | **BD[a]** | **BD[b]** | **BD[c]** | **HD[a]** | **HD[b]** | **HD[c]** | **MT[a]** | **MT[b]** | **MT[c]** | **MM[a]** | **MM[b]** | **MM[c]** |
|---|---|---|---|---|---|---|---|---|---|---|---|---|---|
| Cer d18:1/17:0 | [M+HCOO]- | 1 | 1 | 1 | 1 | 1 | 1 | 1 | 1 | 1 | 1 | 1 | 1 |
| Cer d18:1/17:0 | [M-H]- | 1 | 1 | 1 | 1 | 1 | 1 | 1 | 1 | 1 | 1 | 1 | 1 |
| Cer d20:0/15:0;O(FA 16:0) | [M-H]- | 0.01 | 0.01 | 0.01 | 0 | 0.01 | 0.01 | 0 | 0 | 0 | 0 | 0 | 0.01 |
| FA 12:0 | [M-H]- | 0.81 | 0.52 | 0.85 | 0.92 | 1.04 | 0.79 | 0.01 | 0 | 0.01 | 0.17 | 0.08 | 0.13 |
| FA 14:0 | [M-H]- | 2.41 | 2.66 | 5.63 | 5.21 | 5.06 | 3.38 | 0.15 | 0.07 | 0.21 | 0.18 | 0.3 | 0.12 |
| FA 15:0 | [M-H]- | 0.94 | 1.33 | 2.81 | 1.75 | 2.48 | 1.31 | 0.09 | 0.05 | 0.11 | 0.92 | 1.35 | 1.17 |
| FA 16:0 | [M-H]- | 95.55 | 92.43 | 114.3 | 138.72 | 166.81 | 129.05 | 1.62 | 1.48 | 2.45 | 27.66 | 30.39 | 47.13 |
| FA 16:1 | [M-H]- | 2.57 | 2.91 | 5.11 | 4.07 | 4.6 | 3.05 | 0.24 | 0.12 | 0.49 | 3.41 | 4 | 3.58 |
| FA 16:4 | [M-H]- | 1.32 | 0.97 | 1.24 | 1.16 | 1.27 | 0.99 | 0 | 0 | 0 | 0.04 | 0.1 | 0.21 |

| | | | | | | | | | | | | | |
|---|---|---|---|---|---|---|---|---|---|---|---|---|---|
| FA 17:0 | [M-H]- | 1.26 | 1.62 | 3.3 | 0 | 3.33 | 2.12 | 0.09 | 0.05 | 0.14 | 1.78 | 1.48 | 1.56 |
| FA 18:0 | [M-H]- | 55.86 | 50.45 | 70.69 | 69.44 | 99.09 | 78.84 | 1.36 | 1.61 | 1.63 | 33.16 | 35.45 | 45.19 |
| FA 18:2 | [M-H]- | 0.47 | 0.59 | 0.87 | 0.79 | 0.95 | 0.72 | 0.05 | 0.04 | 0.4 | 1.85 | 1.9 | 1.69 |
| FA 2:0 | [M-H]- | 7.41 | 15.02 | 36.35 | 7.07 | 26.53 | 18.4 | 0 | 0 | 0 | 10.82 | 34.84 | 22.11 |
| FA 20:0 | [M-H]- | 0.86 | 0.84 | 1.53 | 1.04 | 1.78 | 1.45 | 0.02 | 0.02 | 0.04 | 0.43 | 0.24 | 0.5 |
| FA 20:1 | [M-H]- | 0.17 | 0.23 | 0.28 | 0.25 | 0.38 | 0.26 | 0.02 | 0.01 | 0.06 | 0.41 | 0.28 | 0.34 |
| FA 20:5 | [M-H]- | 0.56 | 0.7 | 0.79 | 0.59 | 0.86 | 0.62 | 0.05 | 0.04 | 0.04 | 2.57 | 2.68 | 2.68 |
| FA 22:0 | [M-H]- | 0.35 | 0.48 | 0.86 | 0.49 | 0.98 | 0.64 | 0.02 | 0.01 | 0.04 | 0.18 | 0.16 | 0.22 |
| FA 23:0 | [M-H]- | 0.12 | 0.16 | 0.32 | 0.17 | 0.37 | 0.23 | 0.01 | 0.01 | 0.02 | 0.19 | 0.19 | 0.08 |
| FA 24:0 | [M-H]- | 0.37 | 0.54 | 1.18 | 0.39 | 0.86 | 0.61 | 0.07 | 0.05 | 0.17 | 1.17 | 1.18 | 0.78 |
| FA 26:0 | [M-H]- | 0.25 | 0.35 | 0.67 | 0.29 | 0.53 | 0.42 | 0.04 | 0.02 | 0.1 | 0.56 | 0.38 | 0.47 |
| FA 42:5 | [M-H]- | 0.03 | 0 | 0.44 | 0 | 0 | 0.01 | 0 | 0 | 0 | 0 | 0 | 0 |
| FA 44:7 | [M-H]- | 0.37 | 0.16 | 1.63 | 0.23 | 0.56 | 0.93 | 0.03 | 0 | 0 | 0.59 | 0.28 | 0.34 |
| FA 9:0 | [M-H]- | 0.04 | 0.03 | 0.15 | 0.12 | 0.17 | 0.13 | 0 | 0 | 0 | 0 | 0.11 | 0.16 |
| LNAPE 16:1/N-18:1 | [M-H]- | 0.58 | 0.57 | 0.59 | 0.13 | 0.28 | 0.28 | 0.03 | 0.01 | 0.01 | 0.08 | 0.4 | 0.3 |
| LNAPE 17:0/N-19:0 | [M-H]- | 0.69 | 0 | 3.11 | 0 | 0 | 0 | 0.38 | 0.19 | 0.37 | 8.43 | 10.57 | 8.17 |
| LNAPE 18:1/N-20:1 | [M-H]- | 0.95 | 0.93 | 1.01 | 0.09 | 0.47 | 0.26 | 0.07 | 0.03 | 0.06 | 1.07 | 1.15 | 1.1 |
| LPA 17:1 | [M-H]- | 1 | 1 | 1 | 1 | 1 | 1 | 1 | 1 | 1 | 1 | 1 | 1 |
| LPE 18:0 | [M-H]- | 3.62 | 3.84 | 3.77 | 0.59 | 1.14 | 1.24 | 0.19 | 0.14 | 0.37 | 0 | 0 | 0 |
| LPE 18:1 | [M-H]- | 1.85 | 1.81 | 2.1 | 0.57 | 0.63 | 0.69 | 0.08 | 0.08 | 0.09 | 0 | 0 | 0 |
| LPE 20:4 | [M-H]- | 0.57 | 0.62 | 0.85 | 0.17 | 0.22 | 0.2 | 0.01 | 0 | 0 | 0 | 0 | 0 |

| | | | | | | | | | | | | | |
|---|---|---|---|---|---|---|---|---|---|---|---|---|---|
| LPE 22:6 | [M-H]- | 0.27 | 0.25 | 0.41 | 0.09 | 0.11 | 0.09 | 0.02 | 0.02 | 0.06 | 0 | 0 | 0 |
| LPE O-16:1 | [M-H]- | 0.53 | 0.54 | 0.6 | 0.06 | 0.24 | 0.21 | 0.01 | 0 | 0.11 | 0 | 0.03 | 0.08 |
| LPE O-18:2 | [M-H]- | 0.65 | 0.64 | 0.69 | 0.1 | 0.25 | 0.23 | 0.01 | 0 | 0.04 | 0 | 0.14 | 0.12 |
| LPG 17:0 | [M-H]- | 1 | 1 | 1 | 1 | 1 | 1 | 1 | 1 | 1 | 1 | 1 | 1 |
| LPI 18:0 | [M-H]- | 0 | 0 | 0 | 1.08 | 1.01 | 0 | 0 | 0 | 0 | 0 | 0 | 0 |
| LPS 17:1 | [M-H]- | 1 | 1 | 1 | 1 | 1 | 1 | 1 | 1 | 1 | 1 | 1 | 1 |
| PC 16:0/16:0 | [M+HCOO]- | 0.12 | 0.1 | 0.08 | 0 | 0 | 0.01 | 0.04 | 0.03 | 0.05 | 0.02 | 0.03 | 0.02 |
| PC 16:0/16:1 | [M+HCOO]- | 0.29 | 0.27 | 0.14 | 0.11 | 0.41 | 0.27 | 0.05 | 0.03 | 0.04 | 0.02 | 0.03 | 0.03 |
| PC 17:0/17:0 | [M+HCOO]- | 1 | 1 | 1 | 1 | 1 | 1 | 1 | 1 | 1 | 1 | 1 | 1 |
| PC 17:0/17:0 | [M+CH3COO]- | 1 | 1 | 1 | 1 | 1 | 1 | 1 | 1 | 1 | 1 | 1 | 1 |
| PC 16:0/18:1 | [M+HCOO]- | 0.45 | 0.45 | 0.22 | 0.1 | 0.56 | 0.34 | 0.12 | 0.08 | 0.11 | 0.08 | 0.11 | 0.1 |
| PC 16:0/18:1 | [M+CH3COO]- | 0.57 | 0.54 | 0.29 | 0.18 | 0.71 | 0.43 | 0.15 | 0.11 | 0.15 | 0.1 | 0.14 | 0.12 |
| PC 16:1/18:1 | [M+HCOO]- | 0.31 | 0.28 | 0.15 | 0.19 | 0.49 | 0.31 | 0.06 | 0.03 | 0.04 | 0.02 | 0.03 | 0.03 |
| PC 18:0/18:1 | [M+HCOO]- | 0.21 | 0.21 | 0.09 | 0.09 | 0.29 | 0.19 | 0.06 | 0.04 | 0.06 | 0.04 | 0.05 | 0.05 |
| PC 18:1/18:1 | [M+HCOO]- | 0.27 | 0.26 | 0.14 | 0.08 | 0.38 | 0.22 | 0.05 | 0.04 | 0.05 | 0.03 | 0.05 | 0.04 |
| PC 38:5 | [M+HCOO]- | 0.08 | 0.07 | 0.04 | 0.05 | 0.13 | 0.08 | 0.02 | 0.01 | 0.01 | 0.01 | 0.01 | 0.02 |
| PE 17:0/17:0 | [M-H]- | 1 | 1 | 1 | 1 | 1 | 1 | 1 | 1 | 1 | 1 | 1 | 1 |
| PE 16:0/18:1 | [M-H]- | 0.11 | 0.11 | 0.04 | 0.01 | 0.11 | 0.04 | 0.01 | 0.01 | 0.01 | 0.01 | 0.02 | 0.02 |
| PE 16:1/18:1 | [M-H]- | 0.12 | 0.12 | 0.05 | 0.04 | 0.15 | 0.06 | 0.01 | 0 | 0.01 | 0 | 0.01 | 0.01 |
| PE 18:0/18:1 | [M-H]- | 0.13 | 0.13 | 0.05 | 0.02 | 0.12 | 0.05 | 0.01 | 0.01 | 0.02 | 0.01 | 0.01 | 0.02 |
| PE 18:1/18:1 | [M-H]- | 0.25 | 0.24 | 0.1 | 0.02 | 0.27 | 0.09 | 0.02 | 0.01 | 0.02 | 0.01 | 0.02 | 0.02 |

| | | | | | | | | | | | | | |
|---|---|---|---|---|---|---|---|---|---|---|---|---|---|
| PE 18:0/20:3 | [M-H]- | 0.15 | 0.15 | 0.06 | 0.03 | 0.16 | 0.06 | 0.01 | 0 | 0.01 | 0 | 0.01 | 0.01 |
| PE 18:0/20:4 | [M-H]- | 0.18 | 0.16 | 0.07 | 0.02 | 0.18 | 0.06 | 0.01 | 0.01 | 0.02 | 0.01 | 0.02 | 0.02 |
| PE 18:1/20:4 | [M-H]- | 0.08 | 0.08 | 0.03 | 0.02 | 0.09 | 0.03 | 0.01 | 0 | 0 | 0 | 0.01 | 0.01 |
| PE O-16:1/18:1 | [M-H]- | 0.1 | 0.09 | 0.04 | 0.02 | 0.11 | 0.04 | 0.01 | 0.01 | 0.02 | 0.01 | 0.01 | 0.01 |
| PE O-16:1/20:4 | [M-H]- | 0.13 | 0.14 | 0.05 | 0.04 | 0.18 | 0.06 | 0.02 | 0.02 | 0.02 | 0.03 | 0.03 | 0.04 |
| PE O-18:1/20:4 | [M-H]- | 0.38 | 0 | 0.39 | 0 | 0 | 0 | 0.34 | 0.25 | 0.32 | 0.45 | 0.51 | 0.66 |
| PE O-18:2/20:4 | [M-H]- | 0.14 | 0.14 | 0.06 | 0 | 0.15 | 0.05 | 0.01 | 0.01 | 0.03 | 0.02 | 0.02 | 0.03 |
| PE-Cer 12:1;2O/24:0;O | [M-H]- | 0.11 | 0.1 | 0.05 | 0.02 | 0.05 | 0.05 | 0.01 | 0.01 | 0.01 | 0 | 0 | 0.01 |
| PE-Cer 17:2;2O/24:3 | [M-H]- | 0.75 | 0 | 0.86 | 0 | 0 | 0 | 0.77 | 0.51 | 0.67 | 0.92 | 1.07 | 1.26 |
| PG 17:0/17:0 | [M-H]- | 1 | 1 | 1 | 1 | 1 | 1 | 1 | 1 | 1 | 1 | 1 | 1 |
| PG 16:1/18:1 | [M-H]- | 0.03 | 0.03 | 0.02 | 0.03 | 0.04 | 0.04 | 0.01 | 0 | 0 | 0 | 0 | 0 |
| PG 17:0/19:0 | [M-H]- | 0.01 | 0.01 | 0.01 | 0 | 0 | 0 | 0.01 | 0 | 0.01 | 0 | 0 | 0 |
| PG 18:1/18:1 | [M-H]- | 0.12 | 0.11 | 0.08 | 0.09 | 0.18 | 0.15 | 0.02 | 0.01 | 0.01 | 0 | 0.01 | 0.01 |
| PG 18:1/18:2 | [M-H]- | 0.01 | 0.01 | 0.01 | 0.02 | 0.02 | 0.02 | 0 | 0 | 0 | 0 | 0 | 0 |
| PG 16:1/22:6 | [M-H]- | 0.02 | 0.02 | 0.01 | 0.03 | 0.04 | 0.04 | 0 | 0 | 0 | 0 | 0 | 0 |
| PG 18:1/22:6 | [M-H]- | 0.06 | 0.05 | 0.04 | 0.04 | 0.09 | 0.07 | 0.01 | 0 | 0.01 | 0 | 0.01 | 0.01 |
| PG 22:6/22:6 | [M-H]- | 0.03 | 0.03 | 0.02 | 0.04 | 0.06 | 0.05 | 0 | 0 | 0 | 0 | 0 | 0 |
| PI 18:0/20:3 | [M-H]- | 0.07 | 0.07 | 0.05 | 0.07 | 0.13 | 0.11 | 0.02 | 0.01 | 0.02 | 0.01 | 0.01 | 0.01 |
| PI 18:0/20:4 | [M-H]- | 0.11 | 0.1 | 0.06 | 0.13 | 0.21 | 0.19 | 0.03 | 0.02 | 0.03 | 0.01 | 0.01 | 0.01 |
| PI 18:1/20:3 | [M-H]- | 0.05 | 0.04 | 0.03 | 0.05 | 0.08 | 0.08 | 0.01 | 0.01 | 0.01 | 0 | 0 | 0 |
| PI 18:1/20:4 | [M-H]- | 0.07 | 0.07 | 0.04 | 0.08 | 0.12 | 0.1 | 0.02 | 0.01 | 0.01 | 0 | 0.01 | 0.01 |

| | | | | | | | | | | | | | |
|---|---|---|---|---|---|---|---|---|---|---|---|---|---|
| PI 38:6 | [M-H]- | 0.02 | 0.02 | 0.01 | 0.02 | 0.03 | 0.03 | 0 | 0 | 0 | 0 | 0 | 0 |
| PS 17:0/17:0 | [M-H]- | 1 | 1 | 1 | 1 | 1 | 1 | 1 | 1 | 1 | 1 | 1 | 1 |
| PS 18:0/18:1 | [M-H]- | 0.57 | 0.47 | 0.48 | 0.23 | 0.03 | 0.34 | 0.06 | 0.04 | 0.04 | 0.08 | 0.03 | 0.08 |
| SM d8:1/20:5 | [M+HCOO]- | 0.2 | 0.12 | 0.15 | 0.45 | 0.34 | 0.34 | 0.08 | 0.18 | 0.06 | 0 | 0 | 0.01 |
| SM d18:1/12:0 | [M+HCOO]- | 1 | 1 | 1 | 1 | 1 | 1 | 1 | 1 | 1 | 1 | 1 | 1 |
| SM d18:1/12:0 | [M+CH3COO]- | 1 | 1 | 1 | 1 | 1 | 1 | 1 | 1 | 1 | 1 | 1 | 1 |
| SM 34:1;O2 | [M+HCOO]- | 0.11 | 0.1 | 0.1 | 0 | 0.06 | 0.06 | 0.11 | 0.06 | 0.12 | 0.02 | 0.03 | 0.03 |
| SM 34:1;O2 | [M+CH3COO]- | 0.12 | 0.11 | 0.11 | 0 | 0.06 | 0.06 | 0.1 | 0.07 | 0.14 | 0.02 | 0.03 | 0.03 |
| SM 34:1;O2 | [M+HCOO]- | 0.05 | 0.04 | 0.05 | 0.02 | 0.04 | 0.04 | 0.02 | 0.01 | 0.01 | 0 | 0.01 | 0 |
| SM 42:2;O2 | [M+HCOO]- | 0.07 | 0.06 | 0.06 | 0.02 | 0.05 | 0.04 | 0.04 | 0.03 | 0.05 | 0.02 | 0.02 | 0.03 |
| SMGDG O-18:5/28:7 | [M-H]- | 0.1 | 0.09 | 0.08 | 0.27 | 0.26 | 0.21 | 0 | 0.01 | 0 | 0.01 | 0.03 | 0.04 |
| SMGDG O-19:2/28:7 | [M-H]- | 0.06 | 0.04 | 0.04 | 0.14 | 0.17 | 0.14 | 0 | 0 | 0 | 0 | 0.01 | 0.01 |
| ST 24:1;O4 | [M-H]- | 0 | 0 | 0 | 0 | 0 | 0.01 | 0 | 0 | 0 | 0 | 0 | 0 |
| ST 27:1;O;S | [M-H]- | 0 | 0 | 0 | 0 | 0 | 0 | 0 | 0 | 0.01 | 0 | 0 | 0 |

Abbreviations: BD, chloroform-extraction protocol; MT, MTBE-extraction protocol; MM, MTBE-extraction protocol modified. a, b and c: ratio peak area analyte/ peak area internal standard in triplicate. All these experiments were carried out with 25 seconds of sonication of samples before the extraction protocols. Value = 1 means that are IS samples.

**Supplementary Table 7.** Lipids standards (IS) used for quantification of another class.

| Positive mode | | | Negative mode | | |
|---|---|---|---|---|---|
| **Lipid class*** | **Aduct type** | **IS** | **Lipid class** | **Aduct type** | **IS** |
| Car | [M+H]+ | SM (d18:1/12:0) | FA | [M-H]- | LPA (17:1) |
| PI | [M+H]+ | PC (17:0/17:0) | Lysos | [M-H]- | LPA (17:1) |
| TG | [M+Na]+ | SM (d18:1/12:0) | PI | [M-H]- | PC (17:0/17:0) |
| Cholesterol | [M+Na]+ | SM (d18:1/12:0) | ST | [M-H]- | Cer (d18:1/17:0) |
| NAE | [M+H]+ | Cer (d18:1/17:0) | | | |

***** Lipids class that did not have IS in the same class for use to quantification.

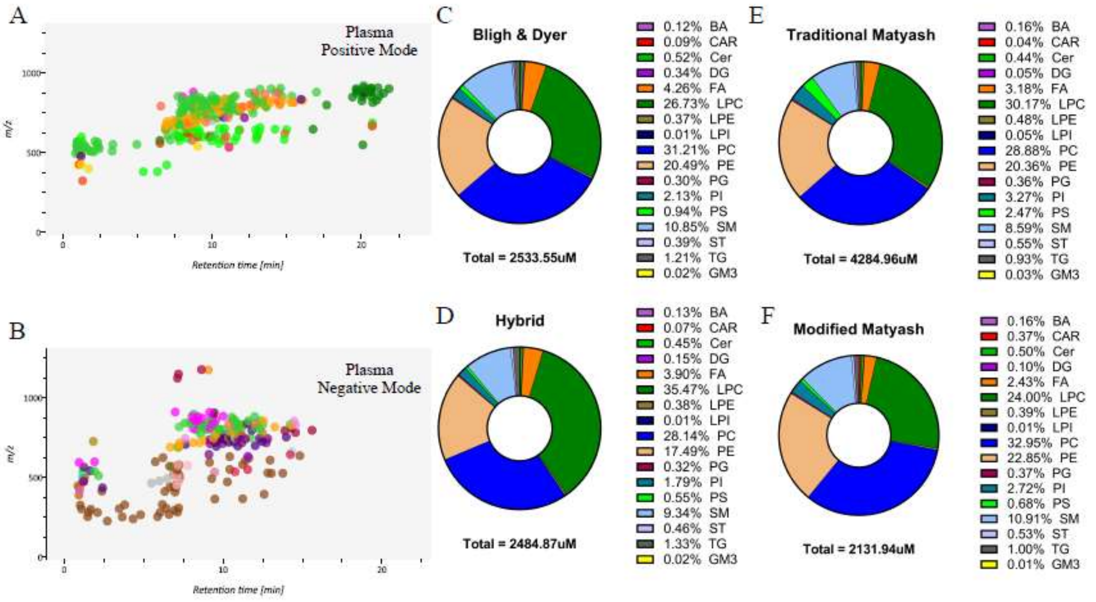


**Fig. S1.** Representative two-dimensional plot (*m/z* vs retention time (RT)) of extracted MS/MS signals and representative features obtained in healthy plasma samples in (A) positive and (B) negative modes acquired by MS-DIAL analysis. Donut plot of the relative composition by lipids subclasses of healthy individual's plasma total lipidome using (C) Chloroform- extraction protocol (BD), (D) Hybrid- extraction protocol (HD), (E) MTBE- extraction protocol (MT), and (F) modified MTBE- extraction protocol (MM) and the total sample quantification.

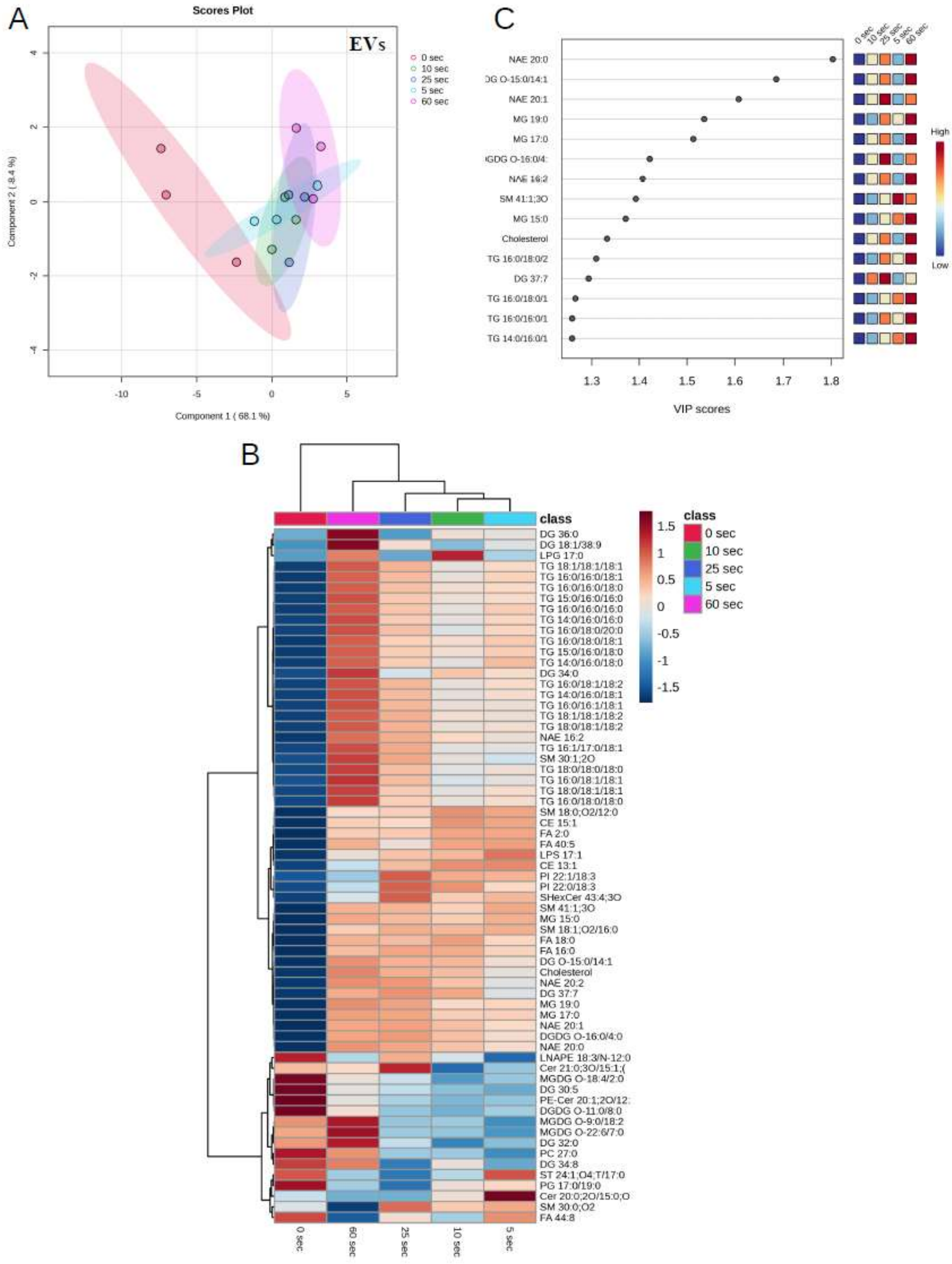


**Fig. S2**. (A) Two-dimensional scatter plot of a partial least square discriminant analysis (PLS-DA) showing individual and technical variation of five different times of EVs sample sonication; 0 seconds (red). 5 seconds (cyan). 10 seconds (green). 25 seconds (blue) and 60 seconds (pink). The colored area represents the 95% confidence interval for each sample group. (B) Hierarchical clustering heatmap based on the relative abundance of lipid species identified in a pool of extracellular vesicles from four different cell phenotypes: M0, M1, M2 and Viral macrophage (MV), in five different times of sample sonication; 0 seconds (red). 5 seconds (cyan). 10 seconds (green). 25 seconds (blue) and 60 seconds (pink). (C) VIP (Variable Importance in Projection) score plot highlighting the 15 most important lipid species for group discrimination based on the PLS-DA.
